# Autocatalytic selection of gene functions in vitro

**DOI:** 10.1101/2025.08.25.672052

**Authors:** Laura Sierra Heras, Christophe Danelon

**Affiliations:** Toulouse Biotechnology Institute (TBI), Université de Toulouse, CNRS, INRAE, INSA, 31077 Toulouse, France

## Abstract

The integration of biological functions into a single operating system is considered a major challenge in the construction of a synthetic cell^1–3^. We present autocatalytic selection (ACS) of gene functions as a driver for integrating biological modules in vitro. A gene of interest (GOI) is introduced into a minimal DNA self-replicator based on the ϕ29 replication machinery^4^ and the function of the GOI is linked to transcription, translation or DNA replication through a positive feedback loop. As the encoded function eventually promotes DNA self-replication, the gene variants with greater activity are selected. Using different coupling mechanisms, we demonstrate ACS of three functions: transcription, synthesis of a deoxynucleoside triphosphate for DNA replication, and β-galactosidase activity. The latter example illustrates how a function that is unrelated to the Central Dogma can be selected. This work paves the way for ACS-driven Darwinian evolution of virtually any biomolecule in vitro, streamlining the construction of increasingly complex synthetic cells as well as the engineering of biotechnologically relevant enzymes.

## Introduction

What does it take to synthesize life? Genome replication, mutation, and selection drive the evolution of new or optimized functions in living cells. This process was probably critical at a certain stage of the origin of life on Earth and represents a milestone in the construction of a synthetic cell in the laboratory^3^. A self-replicating DNA capable of undergoing Darwinian evolution has recently been created^5^. This system involves a two-gene linear DNA template encoding the ϕ29 bacteriophage DNA polymerase (DNAP) and terminal protein (TP), flanked by ϕ29 origins of replication^4^, and could function as a genetic platform for integrating new functionalities. The insertion of genes encoding enzymes for phospholipid synthesis into the self-replicator has recently been demonstrated^6^. However, the combination of the two biological modules—DNA replication and membrane growth—led to a reduced occurrence of active liposomes, revealing the inherent difficulties of module integration. In addition, the design of this genetic circuit does not enable adaptive evolution of the two functions as they are not interconnected^7^. System’s level evolution can serve as a tool for optimizing the individual modules and facilitate their integration^3^. Therefore, an outstanding challenge toward a fully integrated synthetic cell is to implement in vitro surrogate mechanisms for survival-based selection as those found in living organisms.

In in vivo laboratory evolution, the production of a compound of interest is linked to the growth of the strain, allowing the most efficient producers to proliferate^8^. With phage-assisted continuous evolution (PACE), the directed evolution of a protein occurs by linking the desired activity of the protein of interest and phage propagation^9^. By analogy, we propose to couple the activity of a protein or biological module of interest to DNA self-replication in vitro, effectively linking its performance to survival (here the number of DNA molecules) within the system. We termed this mechanism autocatalytic selection (ACS), as selection is driven by the activity of the encoded GOI. By applying selection pressures that favor the desired activity, fitter GOI variants can be enriched through cycles of replication, eventually dominating the population. This method circumvents the constraints of variant screening, for instance the use of an optical output, mimicking the principles of Darwinian selection. A recent commentary reports on a similar concept^10^, suggesting that any gene with a function in the Central Dogma can be integrated and evolved in the minimal self-replicator. We argue that this principle extends beyond genes directly involved in replication, transcription, or translation. As long as a coupling mechanism can be established between the evolvable GOI function and one of the Central Dogma processes, the gene may also be integrated into the replicating template.

Previous studies have shown that the activity of a GOI can be coupled to in vitro translation through the production of amino acids^11–13^. However, this autocatalytic network does not allow for information continuity and Darwinian evolution as DNA replication was not implemented. In compartmentalized self-replication^14^ and related approaches^15–16^, proteins or genetic parts can be evolved by linking their property to the activity of a DNA polymerase and eventually to DNA copy number. These approaches are constrained by in vivo expression and separate amplification of the most active gene variants by polymerase chain reaction (PCR).

In this work, we demonstrate that isothermal ACS enables the integration of functional genetic elements into a DNA self-replicator. We link the GOI’s function to DNA replication through a positive feedback loop, thereby selecting an active gene from a mock library without relying on screening of individual variants. We apply this approach to a transcription-encoding gene and a gene involved in DNA replication. Moreover, as a proof-of-concept to show the general applicability of ACS of gene functions, we select a DNA replicator for β-galactosidase activity. This framework sets the stage for applying in vitro Darwinian evolution to the functional integration of biological modules into synthetic cells.

## Results

### Autocatalytic selection design

We first defined the requirements for the ACS design (Fig. 1a,b):

1. The GOI should be incorporated into the self-replicator seed module. The resulting synthetic DNA therefore consists of the two ϕ29 bacteriophage replication genes coding for the DNA polymerase (DNAP, from gene *p2*) and the terminal protein (TP, from gene *p3*), alongside the GOI, all flanked by ϕ29 origins of replication phosphorylated at the 5’-ends. Instead of a single gene, a multi-gene pathway can be incorporated into the minimal self-replicator as the module of interest.
2. A platform for in vitro transcription-translation-replication (IVTTR) is required. For this purpose, we utilized the PURE system (Protein synthesis Using Recombinant Elements), which provides a minimal and molecularly defined environment for protein expression^17^. To enable self-replication from the expressed proteins, the system is supplemented with replication substrates and cofactors (dNTPs and ammonium sulfate), and two accessory purified proteins (the ϕ29 single-stranded DNA-binding protein or SSB, from gene *p5*, and the double-stranded DNA-binding protein or DSB, from gene *p6*)^4^.
3. To enable functional integration of the GOI, a dependency must be established such that replication occurs only if the GOI is active. This way, only variants producing a functional gene product can amplify, while non-functional variants fail to replicate. The dependency can be introduced at any stage of the Central Dogma—transcription, translation, or replication— through a positive feedback loop, where the activity of the GOI product promotes expression of the replication proteins DNAP and TP, or directly enables replication.
4. To achieve phenotype-genotype coupling, giant liposomes with phospholipid membranes are employed to compartmentalize individual DNA variants. This physical separation restricts the GOI’s function to act on the variant that encodes it, preventing inactive variants from replicating.
5. Establishing the appropriate selection pressure is critical to ensure that replication depends on the activity of the GOI. This involves identifying the conditions that make replication conditional to the GOI’s function.

**Fig. 1.**
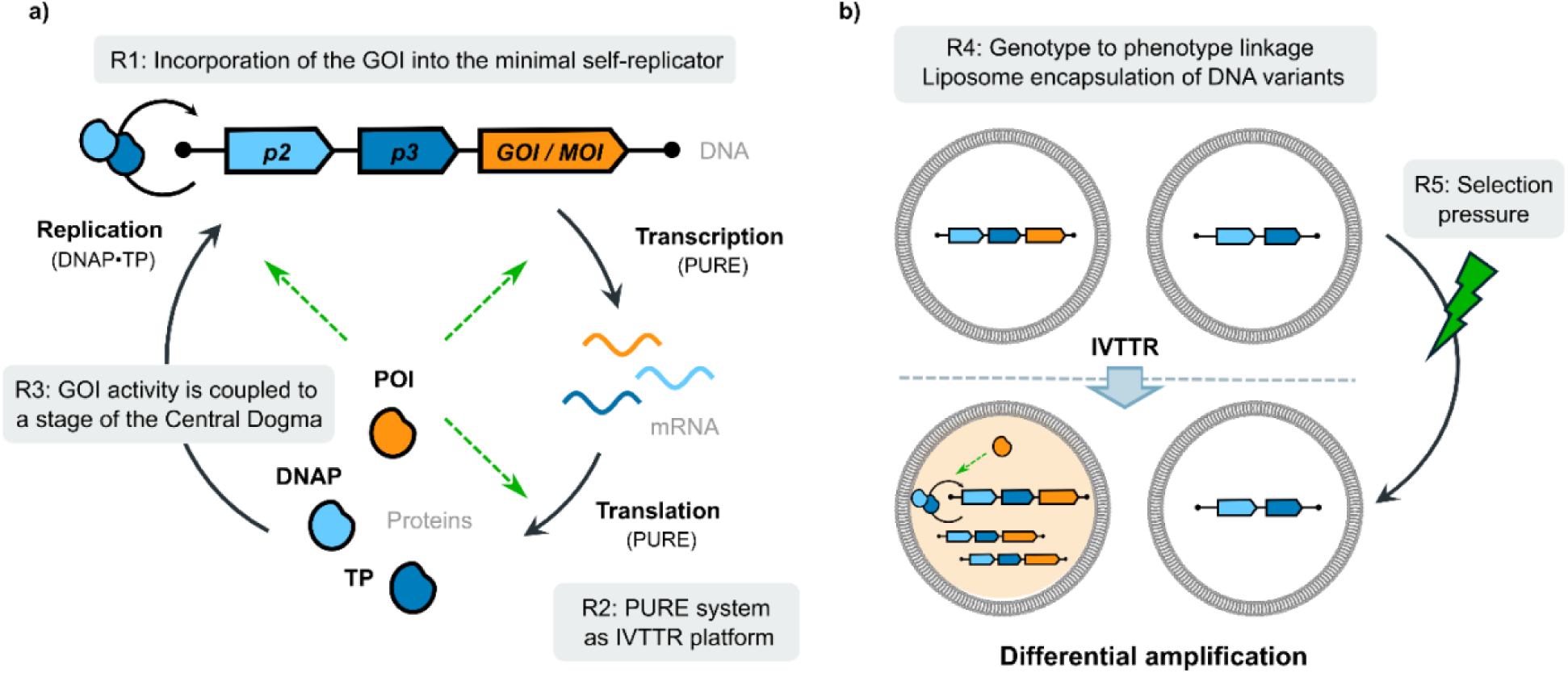
Concept of ACS of gene functions. **a)** Autocatalytic network based on DNA self-replication. Requirement 1 (R1): The gene or genetic module of interest (GOI or MOI) is integrated into the minimal DNA self-replicator. The resulting construct includes the ϕ29 replication genes (*p2* and *p3*), the GOI, and is flanked by the ϕ29 replication origins. R2: Coupled IVTTR is implemented in PURE system supplemented with replication factors. R3: The activity of the GOI is linked to transcription, translation or replication via a positive feedback loop. **b)** Autocatalytic selection of an active gene. R4: DNA variants are encapsulated into liposome compartments for genotype-to-phenotype linkage. This example illustrates a variant encoding the gene of interest (in orange) and another variant lacking that gene. R5: A selection pressure is applied to ensure that replication occurs only if the gene of interest is present/active. This leads to enrichment of the GOI-coding self-replicator. GOI: gene of interest. MOI: module of interest. POI: protein of interest. IVTTR: in vitro transcription-translation-replication.

The process of IVTTR is central to the ACS network. To our surprise, we encountered difficulties in reproducing IVTTR with the minimal self-replicator^5,6^. Further examination indicated that replication was hindered. We hypothesized that the cause could be a change in the formulation of the commercial PURE*frex*2.0. To address this issue, we tested new PURE compositions from separate reagents, guided by the work of Seo and Ichihashi^18^. Through systematic adjustments, we identified Mg²⁺ and NTP concentration as critical factors for successful replication and managed to restore IVTTR efficiency (Fig. S1). We then adopted this modified composition for all experiments involving IVTTR.

Another challenge we encountered was in introducing new genes into the minimal self-replicator using traditional cloning methods. In vitro Gibson assembly or restriction-ligation-based cloning followed by transformation of *E. coli*, repeatedly failed to produce the desired constructs. Plasmid screening by colony PCR or sequencing revealed issues such as mutations in the introduced gene resulting in non-functional variants, incorrect plasmid configurations, or, in cases where the correct plasmid was initially obtained, subsequent growth of the clone led to the loss of the GOI. We hypothesized that these failures originated from leaky expression of toxic proteins during plasmid propagation and/or unintended recombination events in *E. coli* due to repetitive elements, i.e., the regulatory sequences required for expression in the PURE system (namely the T7 promoter, RBS, and T7 terminator). To overcome these issues, we transitioned to yeast as our molecular biology workhorse. This approach involved four key steps: (1) PCR amplification of the genetic elements to be assembled; (2) transformation of the DNA fragments into yeast for in vivo assembly via homologous recombination, (3) isolation of the assembled plasmid, and (4) PCR amplification using 5′ phosphorylated primers to generate linear DNA self-replicators (Fig. S2). This strategy successfully yielded the desired constructs, demonstrating that yeast is a promising candidate for constructing genomes of increasing complexity, facilitating the integration of genetic modules in a synthetic cell.

### Autocatalytic selection of a transcription-encoding DNA self-replicator

To evaluate the potential of the ACS strategy, we first aimed to introduce the gene of the T7 RNA polymerase (T7 RNAP) into the minimal self-replicator. The goal of this design was to create a DNA template capable of self-transcription and replication, which we named *ori-T7-p2p3* (Fig. 2a). Moreover, the integration of T7 RNAP contributes to the broader effort of developing a self-replicating PURE system, where the encoded T7 RNAP autonomously drives transcription—a critical step in regenerating the PURE system itself.

**Fig. 2.**
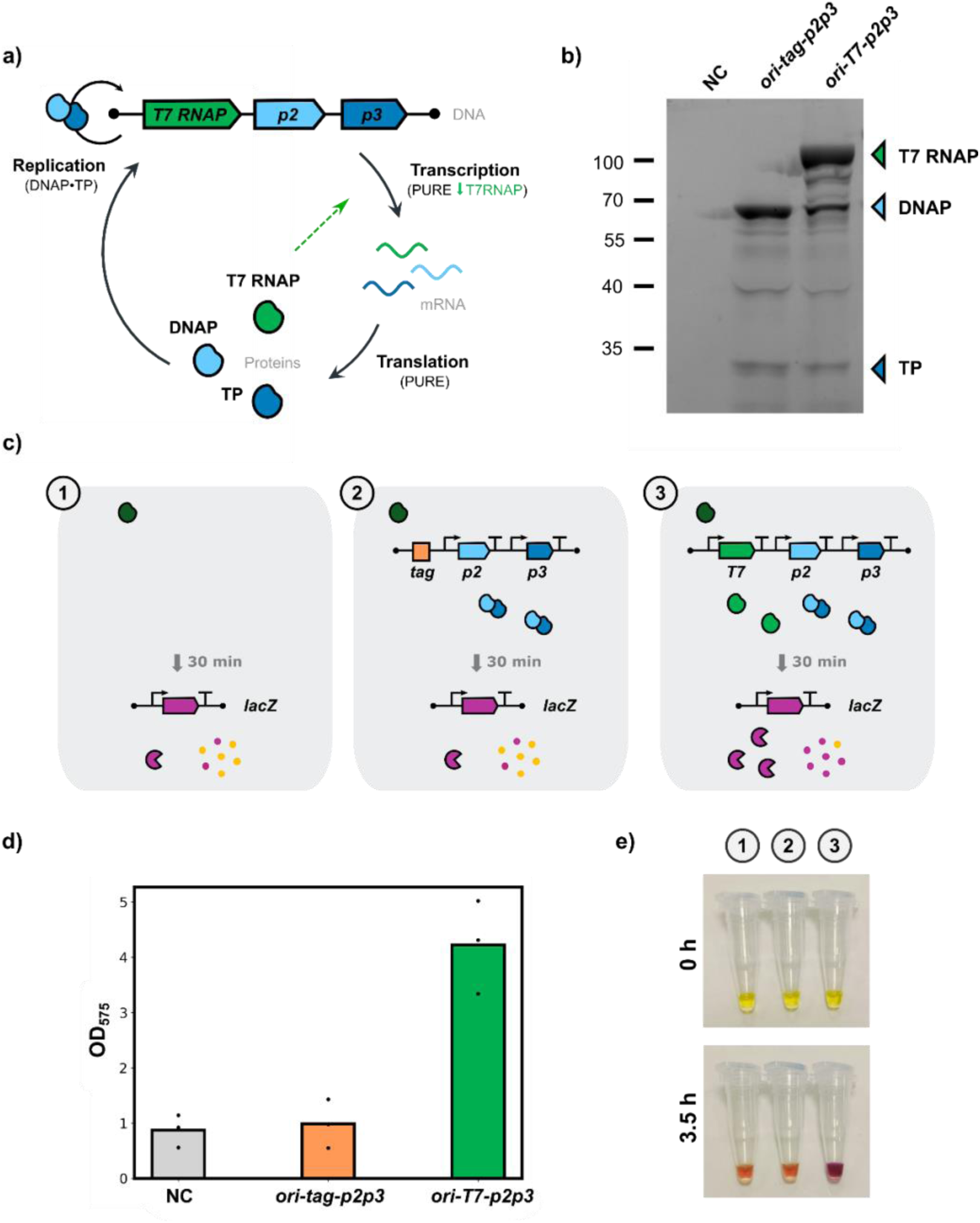
Design and characterization of a DNA template with self-encoded transcription and replication. **a)** Schematic illustration of the synthetic DNA coding for the T7 RNAP, DNAP and TP, flanked by ϕ29 origins of replication. Through a positive feedback loop, the expressed T7 RNAP is expected to rescue transcription that is constrained by the limiting concentration of purified T7 RNAP (selection pressure). This, in turn, enhances the expression of DNAP and TP, driving self-replication. **b)** SDS-PAGE analysis of PURE reaction products using *ori-tag-p2p3* or *ori-T7-p2p3* as DNA template. The PURE solutions were supplemented with GreenLys reagent for specific labelling of the synthesized proteins. NC: negative control with no DNA template. T7 RNAP (∼ 99 kDa), DNAP (∼ 66 kDa) and TP (∼ 31 kDa). **c)** Experimental design to examine T7 RNAP activity. No DNA (1), *ori-tag-p2p3* (2), or *ori-T7-p2p3* (3) was first expressed in the PURE system for 30 min at 37 °C. A *lacZ*- containing DNA template was then spiked into the PURE reaction, along with CPRG to assess β-galactosidase activity by colorimetry. Expression of T7 RNAP from *ori-T7-p2p3* is expected to boost β-galactosidase production. The purified and expressed T7 RNAP are represented in dark and light green, respectively. **d)** Absorbance values at 575 nm after 3.5 h of *lacZ* and CPRG addition. Individual symbols correspond to data from three biological replicates (independent experiments). **e)** Example images of PURE reaction tubes after CPRG addition for the three conditions displayed in c). Images were taken immediately after CPRG addition and 3.5 h later. The conversion of CPRG into CPR, correlating with β-galactosidase expression, is visualized by a color transition from yellow (no/low expression) to orange (moderate expression) and purple (high expression).

By introducing the *T7 RNAP* gene into the DNA template, we intended to establish a positive feedback mechanism that we could use to impose a selection pressure. We hypothesized that under limiting concentrations of externally supplied T7 RNAP, variants encoding a functional T7 RNAP would boost expression, and consequently, gain a replication advantage over non-encoding variants. To test this, we used the minimal self-replicator as the inactive counterpart. Instead of the *T7 RNAP* gene, we included a 130-bp *tag* sequence, *ori-tag-p2p3*, to enable template-specific quantification using quantitative polymerase chain reaction (qPCR). By analyzing the replication profile of the two templates, we sought to identify whether the incorporated functionality conferred a higher fitness under the imposed selection pressure.

As a first step in validating our ACS scheme, we assessed gene expression from the *ori-T7-p2p3* template in PURE system. To visualize the synthesized proteins on gel, fluorescently labelled lysine residues were co-translationally incorporated. Analysis by polyacrylamide gel electrophoresis (PAGE) revealed that all three proteins, T7 RNAP (∼ 99 kDa), DNAP (∼ 66 kDa) and TP (∼ 31 kDa), were produced at detectable levels. Notably, we observed a reduction in the expression levels of *p2* and *p3* compared to the *ori-tag-p2p3* template (Fig. 2b, Fig. S3). This decrease may result from resource sharing as the number of genes increases.

While these results confirm the expression of T7 RNAP, they do not indicate whether the encoded T7 RNAP is active. To this end, we expressed *ori-T7-p2p3* in PURE system for 30 min at 37 °C. Thereafter, we added a second DNA template containing the *lacZ* gene, which encodes the β-galactosidase enzyme. In parallel, we set up two control reactions: one where the *ori-tag-p2p3* construct was pre-expressed before *lacZ* addition, and another one where no DNA was added before spiking the *lacZ* DNA. Our hypothesis was that the increase in T7 RNAP concentration due to expression from the *ori-T7-p2p3* template would boost up the expression of the *lacZ* gene, whereas in both control reactions, expression of *lacZ* would be weaker since it only relies on the purified T7 RNAP provided initially in PURE system (Fig. 2c). To magnify this effect, the concentration of purified T7 RNAP was reduced 100-fold as compared to standard expression conditions. As expected, addition of the β-galactosidase substrate chlorophenol red-β-D-galactopyranoside (CPRG) resulted in a greater colorimetric change in the *ori-T7-p2p3* sample, indicating stronger expression of *lacZ* (Fig. 2d-e, Fig. S4). These results suggest that the T7 RNAP expressed from *ori-T7-p2p3* template is active.

Next, we explored the capability of *ori-T7-p2p3* to drive IVTTR within liposomes. Specifically, we examined how varying the amount of purified T7 RNAP affects the replication output. For comparison, we used the *ori-tag-p2p3* template, whose expression depends solely on the externally supplied T7 RNAP, to assess the impact of T7 RNAP expression from *ori-T7-p2p3*. At standard concentrations of purified T7 RNAP, the *ori-tag-p2p3* template exhibited a greater amplification fold compared to the *ori-T7-p2p3* template, possibly due to the shorter length of the former^6^. However, as the concentration of purified T7 RNAP was reduced, replication of the *ori-tag-p2p3* template dropped, becoming undetectable at 30-fold dilution. In contrast, the *ori-T7-p2p3* template sustained replication even at 100-fold dilution of supplied T7 RNAP (Fig. 3a). These results indicate that, in a regime where purified T7 RNAP is limiting, replication is maintained only when T7 RNAP is regenerated from *ori-T7-p2p3*.

**Fig. 3.**
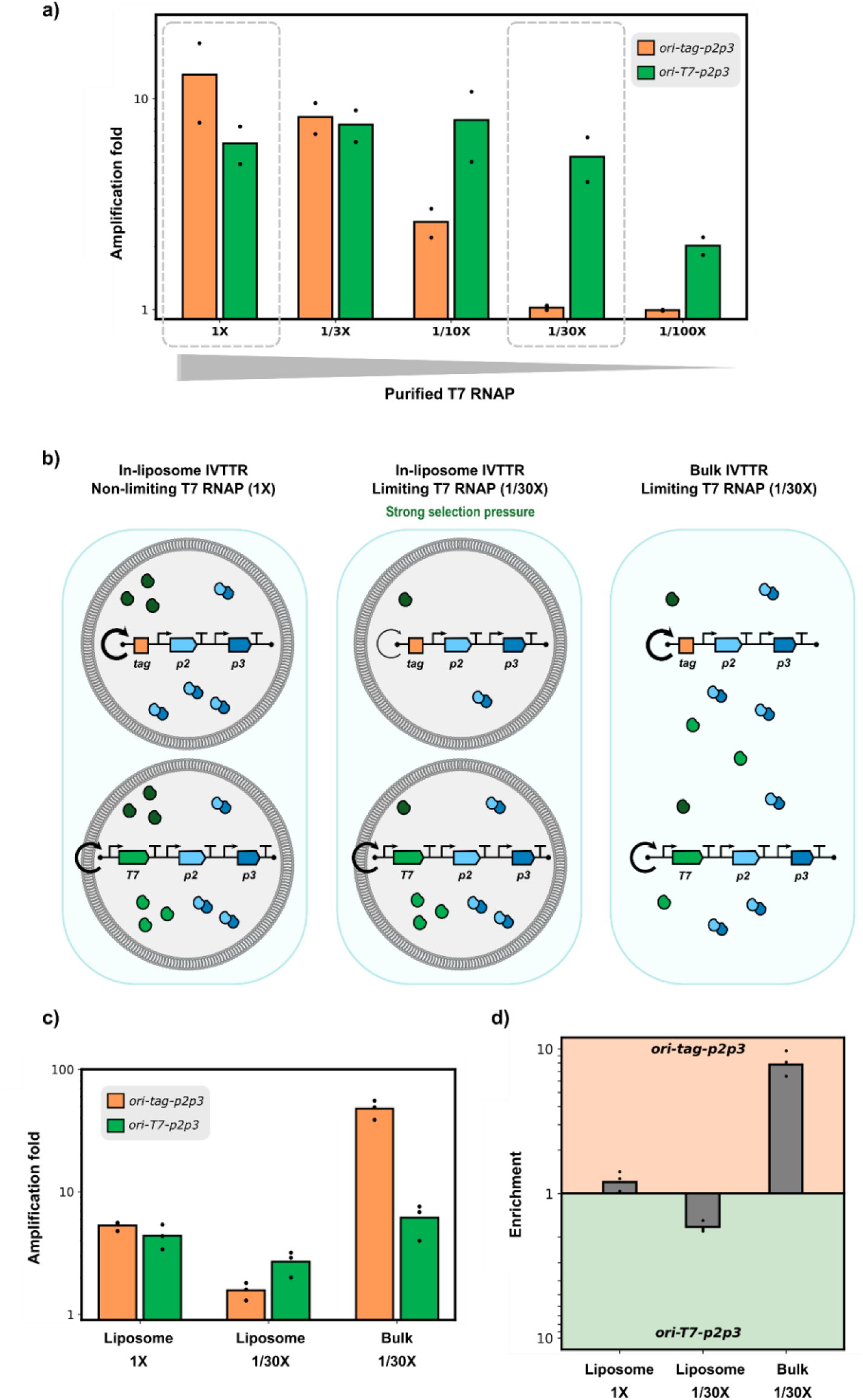
Autocatalytic selection of a transcription-encoding gene. **a)** Amplification folds from in-liposome IVTTR reactions with *ori-tag-p2p3* or *ori-T7-p2p3* as DNA template under different concentrations of purified T7 RNAP. 1X corresponds to the standard T7 RNAP concentration used in PURE system. Quantitative PCR was performed by targeting a region of the *p2* gene. The highlighted conditions (1X and 1/30X) were used in the subsequent competition experiments. Individual symbols correspond to data from two biological replicates (independent experiments). **b)** Schematic illustration of the experimental conditions chosen for selection and the expected outcome. In liposomes under non-limiting purified T7 RNAP concentration (1X), both templates are expected to replicate, with *ori-tag-p2p3* likely being more abundant (bold circular arrow). In liposomes under limiting T7 RNAP concentration (1/30X), *ori-T7-p2p3* is expected to have an advantage since T7 RNAP can be expressed from this template. In bulk under limiting T7 RNAP concentration, both templates are expected to replicate as the produced T7 RNAP is shared in the absence of compartments, with *ori-tag-p2p3* presumably outcompeting *ori-T7-p2p3*. **c)** Amplification folds derived from qPCR data with a 1:1 DNA mixture of *ori-tag-p2p3* and *ori-T7-p2p3* under the three selection conditions explained in b). Total DNA concentration was 500 pM. Quantitative PCR was performed by targeting the *tag* region or the *T7 RNAP* gene. **d)** Enrichments were derived from amplification fold values for the three experimental conditions. Individual symbols correspond to data from three biological replicates (independent experiments).

Building on these findings, we set up a competition experiment between the transcription-encoding and the non-encoding DNA self-replicators under three different conditions (Fig. 3b): (i) In liposomes at standard non-limiting T7 RNAP concentration. Here, we expected both templates to replicate, with the *ori-tag-p2p3* template potentially exhibiting a higher replication efficiency due to its shorter length; (ii) In liposomes at a 30-fold dilution of purified T7 RNAP (strong selection pressure). We anticipated that the *ori-T7-p2p3* template would have a replication advantage, as T7 RNAP can be expressed from the DNA template; and (iii) In bulk at a 30-fold dilution of T7 RNAP. In this case, although the amount of purified T7 RNAP remains restrictive, the lack of compartmentalization would allow the T7 RNAP expressed from *ori-T7-p2p3* to also transcribe *ori-tag-p2p3*. As a result, both templates would replicate, with *ori-tag-p2p3* most likely outcompeting *ori-T7-p2p3*. For the three conditions, the two DNA templates were mixed in a 1:1 ratio as determined by Qubit for a total DNA concentration of 500 pM. The DNA mixture along with the IVTTR reagents were either directly incubated (bulk IVTTR) or encapsulated in liposomes prior to incubation (in-liposome IVTTR). After liposome formation, DNase I was added to prevent replication outside liposomes.

To assess the competition outcome, we performed qPCR targeting the *T7 RNAP* gene and the *tag* region at both the start and end points of the reaction. Amplification folds were calculated for each template (Fig. 3c), and relative enrichments were derived from these values (Fig. 3d). Under a non-limiting T7 RNAP regime in liposomes, the amplification folds for *ori-tag-p2p3* and *ori-T7-p2p3* were 5.3 and 4.4, respectively. This subtle enrichment of the *ori-tag-p2p3* template aligns with observations from individual template experiments, where *ori-tag-p2p3* replicated slightly better than *ori-T7-p2p3* when sufficient T7 RNAP was available (Fig. 3a). Conversely, under limiting T7 RNAP conditions in liposomes, the *ori-T7-p2p3* template achieved an amplification fold of 2.7 compared to 1.6 for *ori-tag-p2p3*, corresponding to a moderate enrichment of 1.7. This enrichment highlights the advantage of *ori-T7-p2p3* when T7 RNAP is restricted. Notably, the *ori-tag-p2p3* variant replicated even though amplification was undetectable at a 30-fold dilution when assessed individually. At the given DNA concentration (250 pM of each template), liposomes may encapsulate more than one DNA copy (see Methods and Figs. S5, S6). Consequently, the two DNA templates can potentially coexist within the same liposome, allowing *ori-tag-p2p3* to replicate to a certain extent due to the T7 RNAP expressed from *ori-T7-p2p3*.

In bulk reactions with a 30-fold dilution of purified T7 RNAP, the amplification folds were markedly different: 47.8 for *ori-tag-p2p3* and 6.1 for *ori-T7-p2p3*. Here, the enrichment for *ori-tag-p2p3* was 6.8, representing a notable amplification advantage. This result reflects the fact that, in bulk, *ori-tag-p2p3* capitalizes on the shared pool of expressed T7 RNAP to the detriment of *ori-T7-p2p3*. In contrast, within liposomes, each template supposedly operates in an isolated environment. The competition outcome is then determined by each template’s capacity to replicate based on its inherent properties, such as length or protein expression levels.

Overall, we demonstrated the feasibility of selecting a transcription-encoding gene from a mixture of replicating DNA. We showed how resource availability and compartmentalization influence the selection dynamics. When T7 RNAP enzyme availability is limited—as opposed to non-limiting conditions—integration of the corresponding gene into the DNA self-replicator provides a distinct advantage within liposome compartments. In contrast, in the absence of boundaries, templates with superior replication properties can seize the shared enzyme pool, leading to a different competition outcome.

### Autocatalytic selection of a metabolism-encoding DNA self-replicator

Next, we aimed to demonstrate the versatility of the ACS design by showing that virtually any GOI can be selected through the engineering of a biomolecular circuit. As a proof of concept, we sought to integrate a metabolism-encoding gene by coupling the activity of the β-galactosidase enzyme to DNA replication, leveraging the well-characterized regulatory mechanism of the *lac* operon. To achieve this, we introduced the *lacZ* gene into the DNA self-replicator, as well as a *lac* operator site upstream of each of the ϕ29 genes. This DNA template was named *ori-Op2Op3-lacZ*. In this network, we hypothesized that the enzymatic conversion of lactose to allolactose by the β-galactosidase enzyme would relieve the transcriptional repression induced by the LacI repressor, enabling DNA self-replication (Fig. 4a). To further examine the system’s dependency on β-galactosidase activity for replication, we introduced a point mutation, E538Q, into the *lacZ* gene, which inhibits the enzymatic production of allolactose^19^. Consequently, with this defective enzyme, derepression would be hindered and DNA replication prevented. The corresponding DNA template was named *ori-Op2Op3-lacZ_E538Q*.

**Fig. 4.**
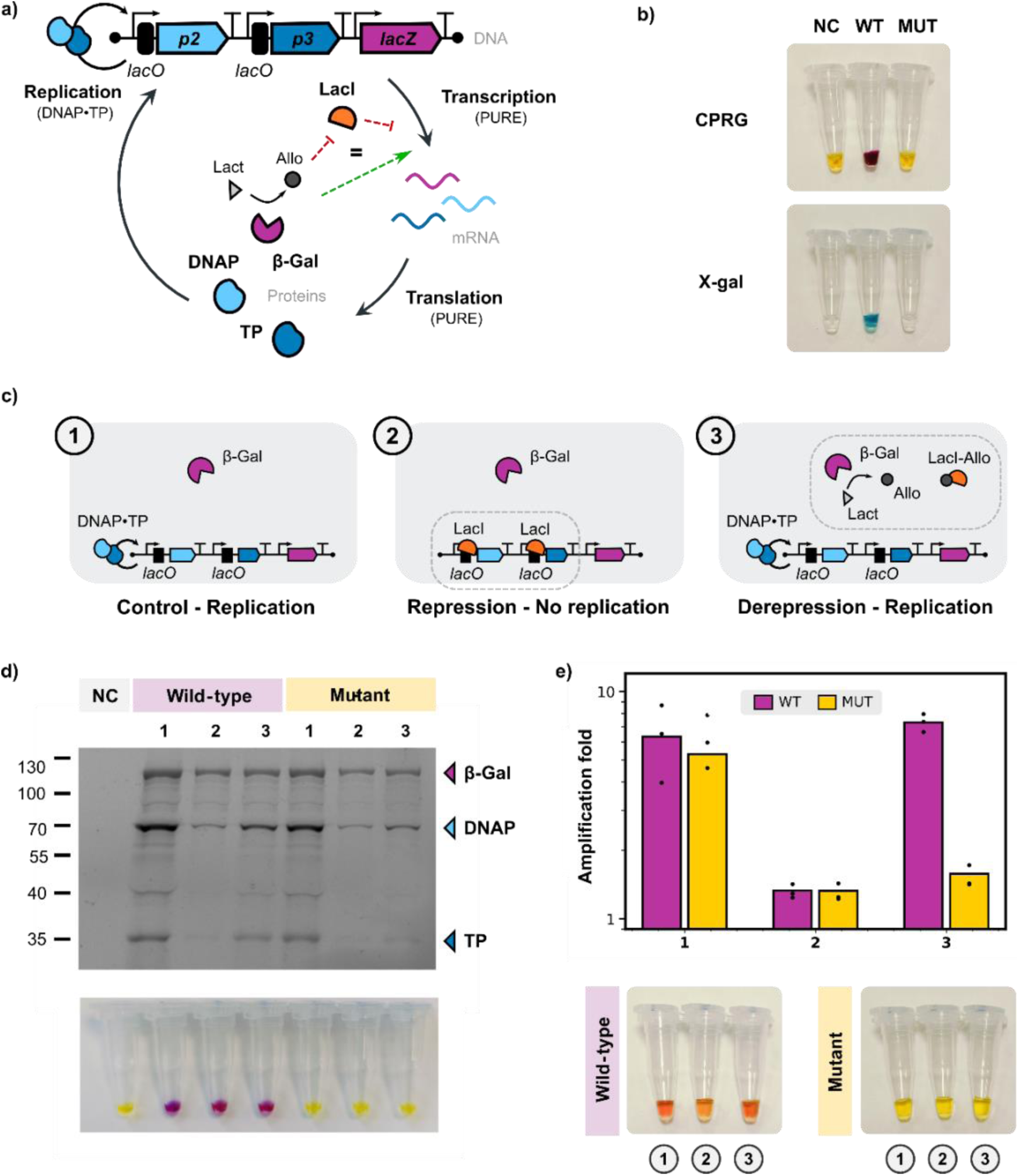
Coupling between the β-galactosidase metabolic module and DNA replication. **a)** Schematic representation of the chimeric DNA self-replicator harboring the ϕ29 replication genes under transcriptional control of *lacO* operator sites and the *lacZ* gene encoding the β-galactosidase. This enzyme converts lactose into allolactose, which is expected to relieve the transcriptional repression exerted by the LacI repressor, enabling expression of the ϕ29 proteins and subsequent template self-replication. **b)** Validation of the metabolic module. PURE reactions after expression of *ori-Op2Op3-lacZ* (WT) or *ori-Op2oP3-lacZ_E538Q* (MUT), and addition of CPRG (top) or X-gal (bottom). Color change in the WT reactions denotes β-galactosidase activity. NC: negative control with no DNA template. **c)** Schematic illustration of the reaction conditions used to validate the coupling between the metabolic and the sensing-replication modules. (1) In the absence of LacI, *p2* and *p3* are expressed allowing replication to occur; (2) In the presence of LacI, transcription is repressed preventing replication; (3) In the presence of LacI and lactose, derepression through β-galactosidase-mediated conversion of lactose into allolactose enables replication. In contrast, if the enzyme is inactive, replication is disabled, as illustrated in (2). **d)** SDS-PAGE analysis of protein expression levels via co-translational labelling under the conditions depicted in c). β-galactosidase (∼ 117 kDa), DNAP (∼ 66 kDa) and TP (∼ 31 kDa). To assay β-galactosidase activity, CPRG was added to the fraction of the PURE reaction that was not used for SDS-PAGE (bottom images). **e)** DNA amplification folds from in-liposome IVTTR reactions with *ori-Op2Op3-lacZ* (WT) or *ori-Op2Op3-lacZ_E538Q* (MUT) under the conditions shown in c). Quantitative PCR was performed by targeting a region of the *p2* gene. Individual symbols correspond to data from three biological replicates (independent experiments). CPRG was added to the liposome samples after IVTTR to indirectly assess DNA replication through β-galactosidase activity (bottom). Increased replication leads to more copies of the *lacZ* gene, which in turn results in higher β-galactosidase expression. A lighter color was observed in the sample containing the wild-type *lacZ* and LacI repressor (WT, sample 2), consistent with a lower replication efficiency. This assay cannot be applied to the mutated template as no β-galactosidase activity is detected, even if replication is enabled.

We first focused on validating the β-galactosidase metabolic module. To do so, we expressed *ori-Op2Op3-lacZ* and *ori-Op2Op3-lacZ_E538Q* in PURE system. After incubation for 1 h at 30 °C, we added CPRG or X-gal, two substrates that enable visual detection of β-galactosidase activity through colorimetric changes. For the sample containing the wild-type *ori-Op2Op3-lacZ* template, addition of CPRG resulted in a color change from yellow to purple. Similarly, with X-gal, the sample transitioned from a transparent solution to a vibrant blue color, confirming enzymatic activity. In contrast, the sample containing the mutant *ori-Op2Op3-lacZ_E538Q* template exhibited no color change with either substrate, remaining yellow and transparent, respectively (Fig. 4b). This confirmed the lack of activity of the β-galactosidase mutant. Moreover, we encapsulated the wild-type *lacZ* as a single-gene DNA template inside liposomes at different concentrations. After CPRG addition and expression for 2 h at 30 °C, we observed an increase in absorbance with higher concentrations of DNA (Fig. S7). These results suggest that CPRG signal may serve as a proxy to qPCR for assessing DNA amplification.

To interrogate the coupling between the metabolic and the sensing-replication modules, we tested the wild-type and mutated DNA templates under three conditions (Fig. 4c): (1) In the absence of LacI repressor, where we expected expression and thereby replication to occur for both DNA templates; (2) In the presence of LacI, where we anticipated transcriptional repression of the *p2* and *p3* genes, thus no replication for either of the two templates; and (3) In the presence of LacI and lactose. Here, we predicted that lactose conversion into allolactose by the wild-type β-galactosidase enzyme would lift transcriptional repression, enabling replication of *ori-Op2Op3-lacZ*. Conversely, the *ori-Op2Op3-lacZ_E538Q* mutant, for which the defective enzyme cannot produce allolactose, was expected not to replicate. Initially, we assessed the system’s behaviour at the level of protein expression (Fig. 4d, Fig. S8). When no repressor was added (1), we observed expression of the three proteins, β-galactosidase (117 kDa), DNAP (66 kDa), and TP (31 kDa), for both templates. Upon addition of LacI (2), expression of the two ϕ29 proteins was significantly diminished for both templates, as anticipated. Interestingly, although there is no *lacO* site upstream the *lacZ* gene, a reduced expression was also observed. This may be due to the presence of an internal *lacO* site within the *lacZ* sequence, which remains subject to partial repression by LacI^20^. When lactose was supplemented to the reaction (3), expression was largely restored for the wild-type template, in line with our expectation that β-galactosidase could convert lactose into allolactose, derepressing gene expression. For the E538Q mutant, a slight derepression was observed, which was unexpected. This effect may be due to a background conversion of lactose into allolactose in PURE system. We routinely observe a slow change of color upon CPRG addition in control samples that do not contain DNA (Fig. S9), suggesting that a β-galactosidase-like side reaction is occurring within the system. Nonetheless, this derepression was less pronounced than upon expression of the wild-type template.

We then evaluated replication dependent on β-galactosidase activity in liposomes under the three conditions described earlier (Fig. 4c). Quantitative PCR data confirmed that the DNA replication outcome was consistent with the protein expression levels (Fig. 4e). In the absence of repressor, both constructs replicated at similar levels. Notably, in the presence of LacI (2), replication was nearly completely prohibited for the two templates. Upon addition of lactose, replication of the wild-type template was restored, whereas the mutated template did not amplify (3), confirming that β-galactosidase-mediated allolactose production is essential for replication. These results demonstrate the successful coupling of the metabolic and sensing-replication modules.

Next, we aimed to perform a competition assay to enrich a metabolically active variant from a mixed DNA population of active and inactive variants. Because differential replication is reliant on genotype to phenotype linkage, we first evaluated whether the allolactose produced by an active variant in one liposome could diffuse and trigger replication in another liposome containing an inactive variant, as this would compromise the selection process. To do so, we prepared two liposome populations: one containing *ori-Op2Op3-lacZ* and the other *ori-Op2Op3-lacZ_E538Q*, both in the presence of LacI and lactose. The populations were mixed in a 1:1 ratio and IVTTR reactions were carried out. If allolactose does not (or poorly) diffuse, only the wild-type template would replicate; if diffusion occurs, both templates would replicate (Fig. S10a). We performed deep sequencing of the liposome-extracted DNA prior and after competition to quantify the fraction of the DNA population corresponding to the wild-type or mutated variant. While both templates were initially present at similar amounts (56% wild-type vs. 44% mutated), the wild-type variant became dominant after IVTTR, representing 80% of the population (Fig. S10b). This result suggests that, if any, allolactose crosstalk between liposomes is minimal and is unlikely to affect the selection process.

We then proceeded with the competition assay to test selective amplification. For this, we mixed the wild-type and mutated DNA templates in a 1:1 ratio and encapsulated the DNA mixture in liposomes together with the IVTTR components. These reactions were conducted in the presence of LacI and lactose (selection pressure), with the expectation that the wild-type template would amplify, while the mutated template would not. As a control to confirm that enrichment depends on β-galactosidase activity being coupled to DNA replication, we performed the same experiment in the absence of LacI and lactose (no selection pressure), where both templates should replicate (Fig. S11a). Confirming this hypothesis, deep sequencing revealed no enrichment without selection pressure (Fig. S11b), even if DNA replication occurred as measured by qPCR (Table S1). Notably, when LacI and lactose were supplied, the metabolically active replicator was successfully enriched, increasing the proportion of the wild-type template in the population from 56% to 72% (Fig. S11b; Table S1). We further studied the system’s ability to enrich an underrepresented, metabolically active self-replicator by mixing the wild-type and mutated DNA templates in a 1:4 ratio (Fig. 5a; Table S1). Again, in the presence of LacI and lactose, the wild-type fraction increased from 31% to 51%, indicating successful selection. No enrichment was observed without selection pressure (Fig. 5b; Table S1).

**Fig. 5.**
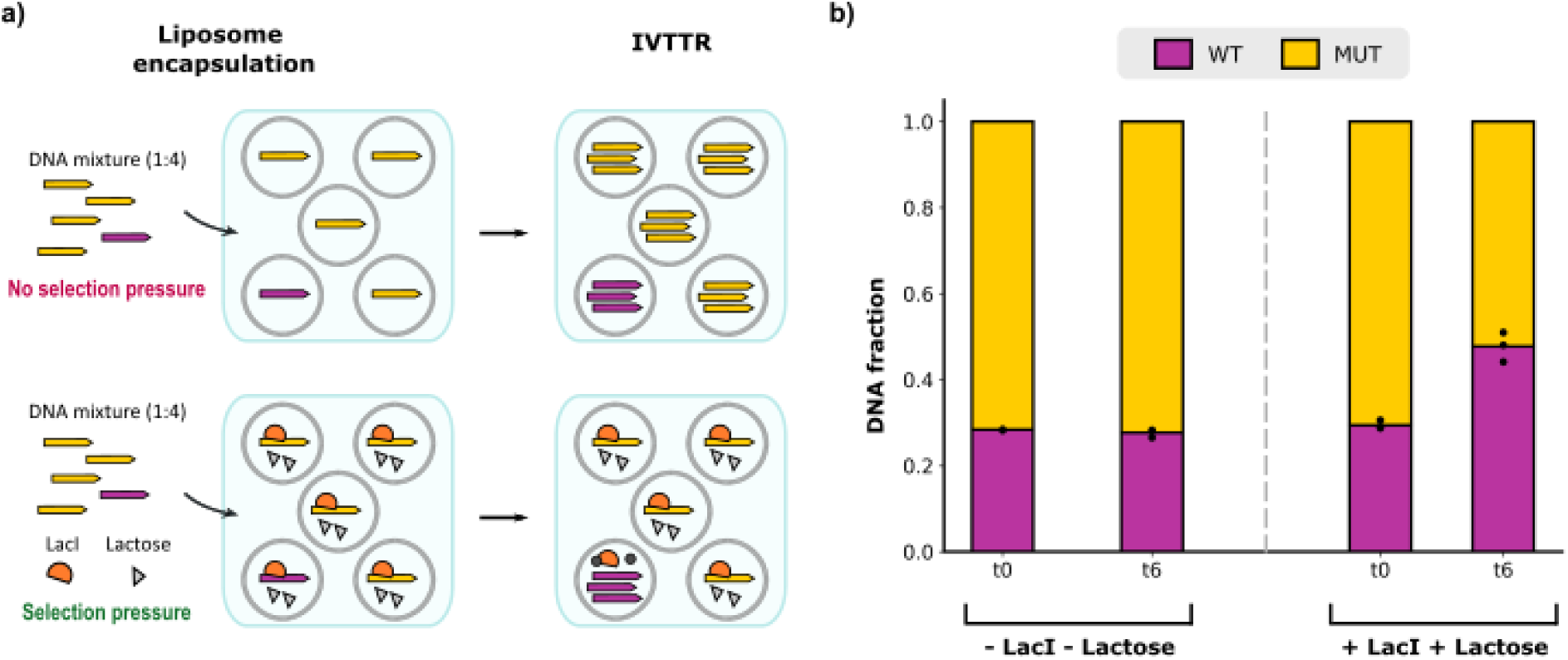
Autocatalytic selection of an active β-galactosidase encoded in the DNA self-replicator. **a)** Schematic illustration of the experimental conditions and predicted replication outcomes. A 1:4 mixture of wild-type (purple) and mutated (yellow) DNA templates was encapsulated along with the IVTTR components into liposomes (total DNA concentration was 500 pM). In the absence of LacI (no selection pressure), both templates are expected to replicate. When LacI and lactose are co-encapsulated (selection pressure), only the wild-type template is expected to be de-repressed providing a replication advantage. **b)** Fraction of the wild-type (WT) and mutated (MUT) variants in the DNA mixture before (t0) and after (t6) IVTTR. The number of reads for each template, as determined by long-read Nanopore sequencing, was used to quantify the relative abundance of each variant. Individual symbols represent data from three biological replicates (independent experiments).

These findings demonstrate that a gene-encoded metabolic function that is arbitrary with respect to transcription-translation-replication can be selected through a simple mechanism that links the desired activity to the activation of DNA replication. Moreover, our findings suggest the potential of this approach for selecting underrepresented, higher-performing variants from a pool of DNA molecules.

### Autocatalytic selection can be driven by a feedback mechanism at the replication level

Thus far, we have shown two different ACS schemes, where the coupling mechanism has been established at the transcriptional level. This approach could, in principle, be adapted to other stages of the Central Dogma, i.e., replication or translation. To explore this idea, we sought to implement a dependency at the level of DNA replication. We introduced the *gmk* gene, which encodes the enzyme guanylate kinase (Gmk), into the minimal self-replicator, yielding the construct named *ori-gmk-p2p3*. Guanylate kinase converts dGMP to dGDP and the enzyme nucleoside diphosphate kinase (Ndk), naturally present in PURE system^17^, subsequently converts dGDP into dGTP, an essential building block for DNA synthesis. Therefore, in a system where dGTP is not supplied externally but dGMP is, the *ori-gmk-p2p3* template catalytically drives its own replication by producing dGTP (Fig. 6a). This creates an autocatalytic selection advantage for the *ori-gmk-p2p3* construct over a control template lacking the *gmk* gene, such as *ori-tag-p2p3*.

**Fig. 6.**
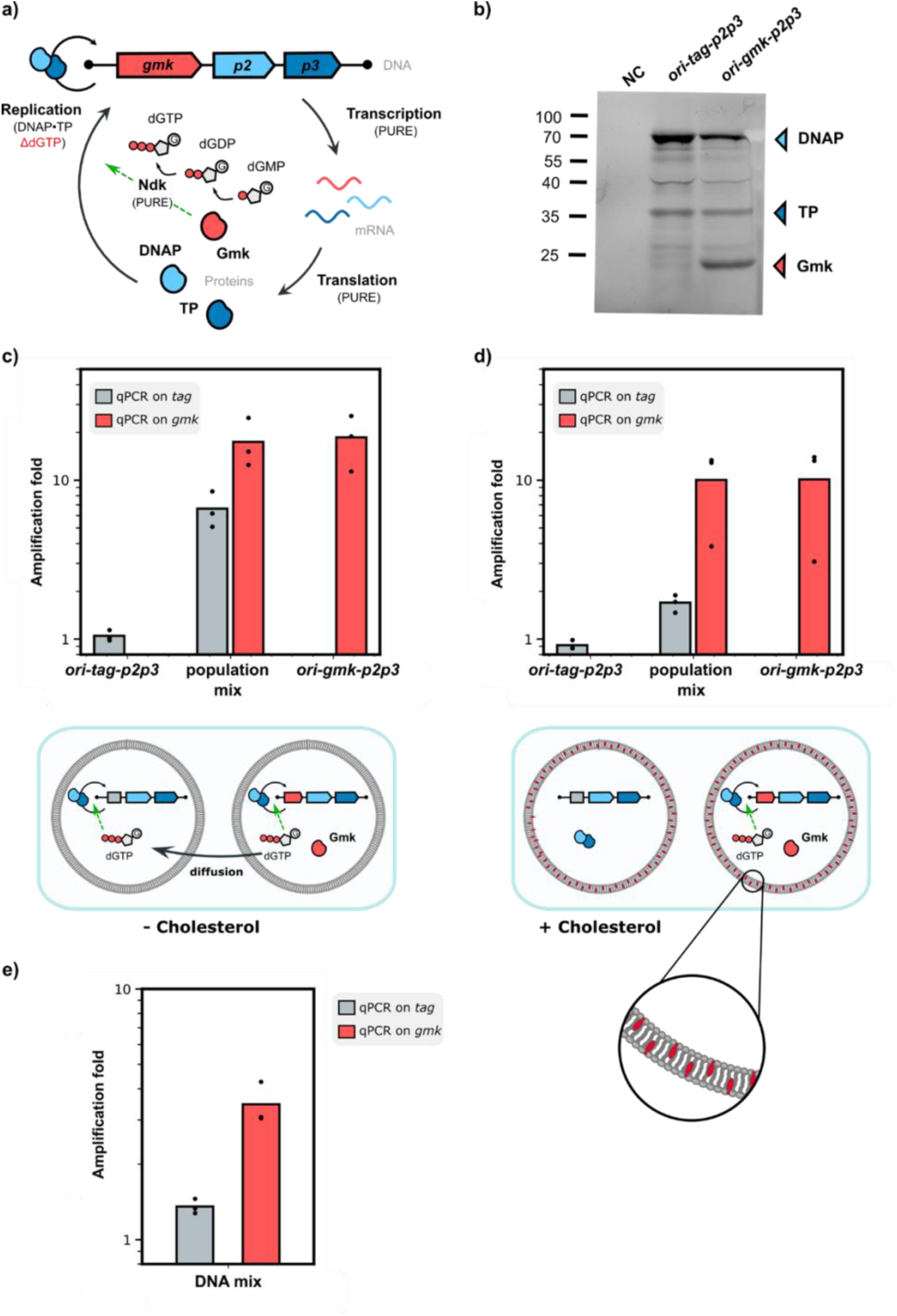
Autocatalytic selection of a dGTP-producing enzyme encoded in the DNA self-replicator. **a**) Schematic representation of the DNA self-replicator including the *gmk* gene. Conversion of dGMP into dGDP by the encoded Gmk and further conversion into dGTP by the PURE-containing enzyme Ndk drives DNA self-replication. **b**) SDS-PAGE analysis of PURE reaction products using *ori-tag-p2p3* or *ori-gmk-p2p3* as DNA template. NC: negative control with no DNA template. DNAP (∼ 66 kDa), TP (∼ 31 kDa), and Gmk (∼ 24 kDa). **c**) Amplification folds from in-liposome IVTTR reactions on dGMP using either *ori-tag-p2p3* or *ori-gmk-p2p3* as the DNA template, or a mix of both liposome populations in a 1:1 ratio. The behavior of the population mix in the dGTP-diffusing and non-diffusing scenarios is illustrated at the bottom of the graph. We suspected that dGTP produced in the *ori-gmk-p2p3* liposomes could diffuse to neighboring *ori-tag-p2p3* liposomes, activating replication. **d**) Same as in c) but with 30% cholesterol-containing lipid membranes. **e**) Amplification folds from in-liposome IVTTR with a 1:1 DNA mixture of *ori-tag-p2p3* and *ori-gmk-p2p3* as DNA template (total DNA concentration was 500 pM). Quantitative PCR was performed by targeting the *tag* region or the *gmk* gene. Individual symbols correspond to data from three biological replicates (independent experiments).

After validating the production of Gmk (∼ 24 kDa), DNAP and TP from the *ori-gmk-p2p3* construct in PURE system (Fig. 6b, Fig. S12), we assessed IVTTR coupled to dGMP conversion. As expected, the *ori-tag-p2p3* construct failed to replicate on dGMP, whereas *ori-gmk-p2p3* exhibited successful amplification, confirming that dGTP can be produced via Gmk-mediated activity, thereby triggering replication (Fig. 8c). However, when mixing the two liposome populations prior incubation, we observed substantial replication of the *ori-tag-p2p3* template (Fig. 8c). We suspected that this effect was due to dGTP diffusion from *ori-gmk-p2p3* to *ori-tag-p2p3-*containing liposomes, inadvertently breaking the genotype-to-phenotype linkage (Fig. 8c). Seeking to mitigate dGTP diffusion, we incorporated cholesterol into the lipid membrane. Cholesterol increases the packing of phospholipids in the liquid-disordered phase, reducing membrane fluidity and permeability to small polar molecules^21^. We repeated the liposome competition assay, this time including 30% cholesterol in the membrane. The differential replication of *ori-gmk-p2p3* over *ori-tag-p2p3* became more pronounced than without cholesterol (5.9 times vs. 2.6-fold). This result suggests that dGTP diffusion was effectively hindered, partially restoring the genotype-to-phenotype linkage. Next, a 1:1 DNA mixture of the *ori-tag-p2p3* and *ori-gmk-p2p3* templates was encapsulated in cholesterol-containing liposomes along with the IVTTR reagents. Quantitative PCR analysis revealed preferential amplification of the *ori-gmk-p2p3* template, indicating successful selection. By leveraging the metabolic coupling between Gmk-mediated dGMP conversion and replication, we have demonstrated that DNA replicators can be engineered to operate when a self-encoded enzyme enables the synthesis of the DNA building blocks, broadening the scope and modularity of our ACS platform.

## Discussion

Selection—the driving force of evolution—acts by favoring traits that enhance survival and propagation. By coupling the activity of a gene of interest to DNA self-replication, we have recreated this natural process in vitro. A distinct ACS mechanism was established for three different gene functions. Through careful scanning of the selection conditions and optimization of the genotype-to-phenotype linkage, we showed that DNA molecules expressing functional gene products outcompete non-functional ones, driving selection of the active variants.

Autocatalytic selection was here applied to a mock library with one active and one inactive variant. However, starting from a larger library for greater sequence diversity using e.g., error-prone PCR, site-saturation mutagenesis with degenerate primers or DNA shuffling, is seamlessly possible. Besides, mutations can be introduced directly inside liposomes during replication by the ϕ29 DNA polymerase, which drives Darwinian evolution of the minimal self-replicator^5^. Selection will arise when mutations in the GOI enhance the fitness of the DNA self-replicator while inactive variants will be purged from the population. Iterative cycles may be required, allowing beneficial mutations to appear and selection to act. This will be particularly useful when selection does not exhibit a high dynamic range or when a gradual increase of the selection stringency is preferred. The evolutionary process may be carried out in a discontinuous mode, involving DNA extraction, PCR, and re-encapsulation, or proceed in a semi-continuous manner^5^, depending on the application. The discontinuous mode allows for precise fine-tuning of component concentrations at each round, whereas the semi-continuous mode may be better suited for developing self-regenerating systems, where the enzyme synthesized in one round is carried over to the next.

We have shown that the feedback mechanism between the GOI function and DNA replication can be established at any of the stages of the Central Dogma—transcription, translation, or replication— offering a wide range of possible functional integrations. Alternative coupling strategies may also be explored, for instance, transporters, ribozymes or allosteric transcription factors other than LacI. Moreover, GOI products that cannot directly act as regulators could be enzymatically transformed into effector molecules for the biosensing and transducing modules^22,23^, thus expanding the repertoire of activating compounds. In addition, logic gates and circuits^24^ can be employed to interconnect different functions, enabling DNA self-replication to proceed based on multimodal inputs. Stepwise implementation of these strategies will further increase the system’s functional information when subjected to the appropriate selection pressure^25^, bringing us closer to the realization of evolving synthetic cells.

Since the biological modules to be integrated may originate from different organisms and have thus not evolved to work together, rounds of in vitro evolution may be necessary to enhance compatibility. Mutations or sequences that resolve conflicts or enhance overall system fitness ensuring the persistence of the DNA self-replicator will be autocatalytically selected, driving evolution toward a more integrated, autonomous system. This gradual buildup of complexity mirrors the natural transition from simple molecular assemblies to functional cellular systems. Autocatalytic chemical cycles—self-sustaining networks of reactions where molecules catalyze their own formation—must have provided a selection advantage in a prebiotic environment, from which key metabolites could have accumulated and core biochemical functions have emerged^26^.

A point of attention for successful ACS implementation is the diffusion of the GOI products and effectors across the liposome membrane, which should be disabled to maintain a strong genotype to phenotype linkage. Molecular diffusion could be impeded by reducing membrane permeability, for instance by adding cholesterol (Fig. 6), by sequestering the permeating molecule or creating a concentration gradient outside liposomes (e.g., allolactose binds to LacI, Fig. 5). Also, it is of practical importance for selection efficiency to carefully choose the concentration of input DNA. In the competition assays involving differentially active DNA variants, individual molecules must be isolated in separate compartments. In the experiments described in Fig. 3, co-encapsulation of *ori-tag-p2p3* and *ori-T7-p2p3* is likely to occur, which may explain why the *ori-tag-p2p3* template was able to replicate (through T7 RNAP produced from co-encapsulated *ori-T7-p2p3*). In fact, at 250 pM DNA concentration, we may even expect a high probability of co-encapsulation (Fig. S6), potentially abolishing selective replication. However, it is important to note that only a fraction of encapsulated DNA may be transcriptionally active^27^. While co-encapsulation may still happen, the probability that both active templates coexist within the same liposome is therefore lower than that calculated from the total DNA amounts, enabling ACS to proceed.

The applicability of ACS extends beyond the construction of synthetic cells. By coupling the activity of the β-galactosidase enzyme to DNA self-replication, we have demonstrated selection of a GOI function that is unrelated to the Central Dogma. This opens the door to leveraging ACS as a driver for in vitro engineering of proteins with a broad range of biotechnological applications. A streamlined *acellular* chassis can help overcome some limitations of in vivo continuous evolution methods, such as metabolic interference, off-target mutagenesis, inadvertent selection pressures and overall controllability.

## Methods

### Construction of DNA fragments

#### General molecular biology techniques

PCRs carried out in this study were performed using the KOD DNA polymerase (MERCK) or the Phusion High-Fidelity DNA Polymerase (Thermo Fisher). For KOD, the reaction mix included 1× Extreme Buffer, 0.02 U µL^−1^ KOD polymerase, ∼0.2 ng µL^−1^ of DNA, 200 nM of each primer (IDT), 0.4 mM dNTPs (Thermo Fisher), and MilliQ water. For Phusion, the reaction mix consisted of 1× Phusion HF Buffer, 0.02 U µL^−1^ Phusion polymerase, ∼0.2 ng µL^−1^ of DNA template, 200 nM of each primer (IDT), 0.2 mM dNTPs (Thermo Fisher), and MilliQ water. When Phusion PCR amplification was unsuccessful due to primer dimer formation, primer concentration was reduced to 20 nM, the solution was supplemented with 5% DMSO, and template concentration was increased to 1 ng µL^−1^. For both KOD and Phusion, the thermocycling protocol was performed according to the manufacturer’s instructions unless stated otherwise. PCR products were purified using the QIAquick PCR Purification Kit (Qiagen).

DNA fragments were in vitro assembled using the NEBuilder HiFi DNA Assembly system (NEB), and the resulting constructs were transformed into chemically competent TOP10 *E. coli* cells prepared in-house. Alternatively, 2 µL of the NEBuilder HiFi DNA Assembly reaction were directly used as template for PCR amplification. Diagnostic colony PCR was carried out using the 2x Phire Green HS II Master Mix (Thermo Fisher) according to the manufacturer’s protocol except that the volume was downscaled to 10 µL. Plasmid isolation from *E. coli* was performed using the EZ-10 Spin column plasmid DNA miniprep kit (Bio Basic). Sequence verification was performed using either Sanger sequencing (Eurofins Genomics) or long-read Oxford Nanopore technology (Plasmidsaurus).

Primers used for PCR and qPCR are listed in Tables S2 and S3, respectively. All plasmids used and constructed in this study are listed in Table S4.

#### ori-T7-p2p3

To clone the T7 RNA polymerase (T7 RNAP) gene under control of T7 regulatory sequences, the *T7 RNAP* gene was PCR-amplified from plasmid G442 using primers ChDT-1365 and 1366, while plasmid G435 (G340 from Abil et al. ^5^ with a synonymous mutation in *p2* to remove a *PmeI* restriction site) was linearized with primers ChDT-1363 and 1364. Both fragments were assembled in vitro using 20 fmol of each in a 10 µL NEBuilder HiFi DNA Assembly reaction. Transformation of *E. coli* with the assembly product always led to plasmids with incorrect sequences. Therefore, we switched to the yeast *Saccharomyces cerevisiae* (strain CEN.PK2-1C) as our assembly platform.

Plasmid pY002, containing *ori-T7-p2p3*, was in-yeast assembled from three DNA fragments: (1) *oriL-T7RNAP*, amplified from the assembly product generated as described above using primers *ChDT-41* and *42; (2) p2p3-oriR*, amplified from G435 with primers *ChDT-43* and *44 and (3) CEN6/ARS4-URA3*, comprising the yeast centromeric replication origin, *CEN6/ARS4*, and the auxotrophic marker *URA3* for maintenance and selection in yeast, respectively. This fragment was amplified from pRS316 with primers ChDT-56 and 57. To enable efficient assembly in yeast, PCR primers were designed to produce fragments with 60-bp overlaps. The DNA fragments were mixed at a concentration of 100 fmol for the *CEN6/ARS4-URA3* fragment and 200 fmol for the other two DNA fragments. Transformation in *S. cerevisiae* CEN.PK2-1C was performed using the lithium acetate/single-stranded carrier DNA/polyethylene glycol method^28^. For selective growth, cells were plated on complete supplemental medium without uracil (CSM-URA) agar plates consisting of 6.7 g L^−1^ yeast nitrogen base without amino acids (Formedium), 20 g L^−1^ glucose (Formedium), 0.77 g L^−1^ CSM-URA (MP Biomedicals), and 20 g L^−1^ agar (Euromedex). Incubation was carried out at 28 °C for 2 days. Plasmid extraction from yeast was performed as previously described^6^. Afterwards, 1 µL of the extracted DNA was used in a PCR reaction with primers ChDT-491E and 492E to generate *ori-T7-p2p3*. The resulting linear amplicon was sequence-verified using long-read Nanopore technology.

#### ori-Op2Op3-lacZ

Plasmid pT002, containing two *lacO* sites located after the T7 promoter regions of both *p2* and *p3* genes, was derived from plasmid G435. To introduce the *lacO* sequence upstream *p2*, G435 was amplified with primers ChDT-1 and 2, which contain an overhang with the *lacO* sequence. The amplicon was then recircularized though in vitro assembly and transformed into *E. coli*. After plasmid isolation, the resulting plasmid, pT001, was used as a template to introduce the second *lacO* site upstream the *p3* gene. To this end, pT001 was linearized using primers ChDT-19 and 20. Subsequently, a double-stranded DNA (dsDNA) molecule containing the *lacO* sequence and overhangs complementary to the linearized pT001 backbone was generated by annealing two complementary single-stranded DNA (ssDNA) primers, ChDT-21 and 22. The ssDNA primers were mixed at a concentration of 100 nM each, heated to 95 °C for 5 min, and gradually cooled down to 25 °C over the course of 1 h. The formation of the annealed dsDNA was confirmed on a 2.5% agarose gel. The dsDNA was then assembled into the linearized pT001 backbone to generate pT002, which was sequence-verified by Sanger sequencing.

Plasmid pT003, which contains the *lacZ* gene under control of a T7 promoter and terminator, was generated via in vitro DNA assembly. The T7 regulatory sequences were taken from plasmid G435 by PCR amplification with primers ChDT-7 and 8. The *lacZ* gene was amplified from pBADMyc-HIS-lacZ using primers ChDT-11 and 12, which introduced a start and stop codon. This amplified product served as a template for a second PCR reaction with primers ChDT-9 and 10, which incorporate overhangs for assembly into the linearized G435 backbone. After transformation and isolation from *E. coli*, the resulting plasmid was sequence-verified by Sanger sequencing.

Again, due to difficulties to generate the plasmid containing *ori-Op2Op3-lacZ* in *E. coli*, we continued the assembly in yeast. *Saccharomyces cerevisiae* was co-transformed with three DNA fragments to generate plasmid *pY002:* (1) *oriL-Op2Op3*, amplified from pT002 with primers ChDT-41 and 47; (2) *lacZ-oriR*, amplified from pT003 with primers ChDT-44 and 48; and (3) *CEN6/ARS4-URA3* fragment from pRS316 with primers ChDT-56 and 57. Extraction from yeast, production of the linear *ori-Op2Op3-lacZ* via PCR and sequence verification by long-read Nanopore were performed as described above.

#### ori-Op2Op3-lacZ_E538Q

To generate the construct *ori-Op2Op3-lacZ_E538Q*, a site-specific mutation was introduced in the *lacZ* gene, where <u>G</u>AA (encoding glutamic acid, E) was mutated to <u>C</u>AA (encoding glutamine, Q) at position 538, resulting in the E538Q substitution. This mutation was introduced by amplifying plasmid pT003 using primers ChDT-72 and 73. The mutated plasmid was then recircularized by in vitro assembly. Two microliters of the assembled product were used as a template in a PCR reaction with primers ChDT-44 and 48 to generate *lacZ_E538Q-oriR*. This fragment, along with the *oriL-Op2-Op3* and *CEN6/ARS4-URA3* fragments, were co-transformed into yeast to construct plasmid pY008. In this case, the *CEN6/ARS4-URA3* fragment was amplified from pRS316 with primers ChDT-57 and 65 to incorporate an ampicillin resistance gene and an *E. coli* replication origin to facilitate future modifications in *E. coli* when needed. This fragment was then renamed *ColE1-Amp-CEN6/ARS4-URA3*. Extraction from yeast and production of the linear *ori-Op2Op3-lacZ_E538Q* for IVTTR was performed as described above. Sequencing of the linear PCR product using long-read Nanopore technology confirmed the presence of the E538Q (<u>G</u>AA > <u>C</u>AA) mutation. Three additional mutations were found in the *lacZ* gene R60H (C<u>G</u>C > C<u>A</u>C), Y101H (<u>T</u>AC > <u>C</u>AC), and Y106H (<u>T</u>AT > <u>C</u>AT).

#### ori-gmk-p2p3

The construction of *ori-gmk-p2p3* was carried out in a similar manner to that of *ori-T7-p2p3*. The *gmk* gene was PCR-amplified from *E. coli* BW25113 genomic DNA using primers ChDT-199 and 200. The *gmk* amplicon was then assembled in vitro into G435 linearized with primers ChD-1363 and 1364. This assembly product served as a template to generate the *oriL-gmk* fragment by PCR using primers ChDT-41 and 42. For in vivo assembly, yeast was co-transformed with *oriL-gmk*, *p2-p3-oriR*, and *ColE1-Amp-CEN6/ARS4-URA3* to generate plasmid pY011. Extraction from yeast, production of the linear *ori-gmk-p2p3* and sequence verification were performed as described above. A synonymous mutation was found in the *p2* gene K179 (AA<u>A</u> > AA<u>G</u>).

#### ori-tag-p2p3

To introduce a *tag* sequence into the *ori-p2p3* minimal self-replicator, enabling selective qPCR against *ori-T7-p2p3* and *ori-gmk-p2p3*, a 170-bp *tag* was first amplified from plasmid pRS316 using primers ChDT-079 and 080. Plasmid G435 was then linearized with primers ChDT-077 and 078. The linearized G435 plasmid and the *tag* amplicon were assembled in vitro, and the assembly product was used for *E. coli* transformation. Correct insertion of the *tag* sequence was confirmed by colony PCR. After plasmid isolation (pT007), the linear *ori-tag-p2p3* template for IVTTR was generated as described above.

### Fluorescence labelling of in vitro synthesized proteins and gel analysis

Standard 10 µL PURE*frex*2.0 (GeneFrontier Corp.) reaction mixtures containing 5 µL of Solution I, 0.5 µL of Solution II, 1 µL of Solution III, 0.5 U μL^−1^ of Superase·In RNase inhibitor (Thermo Fisher), and 0.5 nM of linear DNA template were supplemented with 0.25 µL of BODIPY-Lys-tRNA_Lys_ (FluoroTect Green_Lys_, Promega) to fluorescently label lysine residues in the proteins being synthesized. When required, 350 nM of purified LacI repressor and 2 mM of lactose were added to the mixture. Protein expression was carried out at 30 °C for 4 h and samples were treated with 0.3 mg mL^−1^ RNase A (Thermo Fisher) at 37 °C for 1 h 30 min. Next, proteins were denatured in NuPAGE LDS Sample Buffer (Thermo Fisher) and 10 mM of dithiothreitol (DTT, Formedium) at 65 °C for 5 min. Ten microliters of the mixture were resolved on an in-house prepared 12% SDS polyacrylamide gel electrophoresis (SDS-PAGE) gel. The SDS-PAGE gels were run at 100 V for 20 min followed by 150 V for 40 min. Detection of fluorescently labelled translation products was carried out using a fluorescence gel imager (Bio-Rad ChemiDoc Imager).

### T7 RNAP activity assay

Standard PURE*frex*2.0 reactions were prepared by mixing 5 µL of Solution I, 0.5 µL of Solution II without T7 RNAP, 1 µL of 100-fold diluted T7 RNAP (GeneFrontier Corp.), 0.25 µL of Solution III, 0.25 µL of 20 U µL^−1^ Superase·In RNase inhibitor, 0.65 µL of nuclease-free water, and 0.5 µL of either 10 nM *ori-tag-p2p3*, 10 nM *ori-T7-p2p3*, or nuclease-free water. After pre-incubation at 30 °C for 30 min, 0.5 µL of 10 nM *lacZ* template (0.5 nM final concentration) and 0.6 µL of 10 mg mL^−1^ Chlorophenol Red-β-D-galactopyranoside (CPRG, MERCK) were added. The *lacZ* transcriptional unit was amplified from pT003 using primers ChDT-16 and 17. Pictures were taken immediately after CPRG addition and following 3.5 h of incubation at 30 °C (4 h including pre-incubation). Additionally, after 3.5 h, samples were diluted 10-fold in Milli-Q water, and optical density was measured at 575 nm (OD_575_) for chlorophenol red (CPR) detection using a PowerWave Select X Microplate Reader (Bio-Tek Instruments) and a 384-well, clear-bottom, black microplate (Greiner BIO-ONE).

### Optimization of PURE composition for IVTTR reactions

Due to challenges encountered in reproducing replication in standard PURE*frex*2.0-based IVTTR reactions^5,6^, we tested different PURE Solution I compositions assembled from individual components. The two modified compositions, modified 1 (M1) and modified 2 (M2), that helped us identify the key components affecting replication are reported in (Fig. S1b). As M1 successfully restored replication (Fig. S1), it was used in all IVTTR experiments described in this work. A regular 20-µL IVTTR reaction using PURE*frex*-M1 consisted of 7.24 µL of Solution I, 4.2 µL of Solution II, 1.33 µL of Solution III, 20 mM of ammonium sulfate, 600 µM of dNTPs (Promega), 375 µg mL^−1^ of SSB protein and 105 µg mL^−1^ of DSB protein (purified as described by van Nies et al. ^4^), 0.5 nM of DNA, and nuclease-free water (Thermo Fisher). DNA templates were measured with Qubit 2.0 Fluorometer (Thermo Fisher) using the Qubit dsDNA Broad Range Assay kit (Invitrogen).

### IVTTR reactions with gene-encoded T7 RNAP transcription

IVTTR reactions were assembled using PURE*frex*-M1 as described above, except that 3.6 µL of Solution II without T7RNAP was used, and 0.6 µL of T7 RNAP was supplemented at the desired concentration. T7 RNAP dilutions (3, 10, 30, or 100-fold) were prepared using Solution II dilution buffer (GeneFrontier Corp.). Dilutions were freshly prepared before use as we observed that the diluted T7 RNAP loses activity over time. DNA templates consisted of either 0.5 nM of *ori-tag-p2p3* or *ori-T7-p2p3* for individual template experiments, or a 1:1 mixture (0.25 nM of each) for competition experiments.

For bulk IVTTR, reactions were incubated directly in nuclease-free PCR tubes at 30 °C for 16 h. To assess the level of amplification, samples were collected prior to (start-point sample) and following (end-point sample) incubation. For in-liposome IVTTR, reactions were assembled in 1.5 mL Eppendorf tubes and 10 mg of pre-desiccated lipid-coated beads were added to the reaction. Lipid-coated beads composed of DOPC, DOPE, DOPG and cardiolipin (50:36:12:2 molar ratio) were prepared as described by Abil et al.^5^, excluding DHPE-Texas Red. The tubes were placed in an automatic tube rotator (VWR) and rotated along their axis for 1 h at 4 °C. Subsequently, four freeze-thaw cycles were applied to the samples. Using a cut pipette tip to widen the aperture and prevent liposome disruption, 12.5 µL of the liposome suspension was transferred to a nuclease-free PCR tube containing 1.5 U of DNase I (NEB). Reactions were incubated at 30 °C for 20 min to allow DNase I activity. Afterward, a sample was taken (start-point sample), and the incubation was continued for up to 16 h (end-point sample). DNase I was inactivated by heating at 75 °C for 15 min. Both liposome and bulk reaction solutions were diluted 100-fold in Milli-Q water prior qPCR analysis.

### Quantitative PCR

Quantitative PCR reactions consisted of 1× PowerUP SYBR Green Master Mix (Applied Biosystems), 400 nM of each primer (Table S3), 1 μL of 100-fold diluted sample, and Milli-Q water to a final volume of 10 μL. The thermal cycling program and data acquisition were performed on a Quantstudio 3 Real-Time PCR system (Thermo Fisher), utilizing a cycling protocol of 2 min at 50 °C, 5 min at 94 °C, (15 sec at 94 °C, 15 sec at 56 °C, 30 sec at 68 °C) × 40 cycles, ending with a melting curve from 65 °C to 95 °C. Data analysis was conducted with the Quantstudio Design and Analysis software v1.5.2 (Thermo Fisher). Amplification folds were calculated using the Δ*Ct* method, corrected for primer efficiency (*E*), using the formula: *E*^Δ*Ct*^. Primer efficiency was determined from a standard curve generated using serial 10-fold dilutions of the corresponding DNA templates, with concentrations ranging from 1 fM to 1 nM. Efficiency was then calculated using the formula: *E* = 1^−1/*m*^, where *m* is the slope of the standard curve. Relative enrichments were calculated using the Pfaffl method, i.e., the 2^−ΔΔ*Ct*^ method with efficiency correction: *E*_target1_^Δ*Ct*-target1^ / *E* ^Δ*Ct*-target2^. Target 1 refers to the *tag* region and target 2 the *T7 RNAP* or *gmk* region.

### β-Galactosidase activity assay

Standard 10 µL PURE*frex*2.0 reactions consisting of 5 µL of Solution I, 0.5 µL of Solution II, 0.25 µL of Solution III, and either 0.5 nM of linear DNA template (*ori-Op2Op3-lacZ* or *ori-Op2Op3-lacZ_E538Q*) or water, were incubated at 30 °C for 1 h. Then, 2 µL of CPRG (10 mg mL^−1^) or X-gal (200 mg mL^−1^) were added. Images were taken directly after substrate addition for CPRG and 30 min later for X-gal to allow for product precipitation.

To assess β-Galactosidase activity in liposomes, 20 µL PURE*frex*2.0 reactions at varying concentrations of DNA, 0 pM, 50 pM 100 pM, 500 pM and 1000 pM, were assembled on ice. Liposomes were prepared as described above and were treated with 1.5 U of DNase I to avoid expression outside liposomes. CPRG was added to the samples and liposomes were incubated at 30 °C for 2 h. After incubation, pictures were taken and OD_575_ was measured in a PowerWave Select X Microplate Reader (Bio-Tek Instruments) using a 384-well, clear-bottom, black microplate (Greiner BIO-ONE).

### IVTTR reactions with gene-encoded β-Galactosidase

IVTTR reactions were prepared using PURE*frex*-M1 as described above. When indicated, 350 nM of LacI and 2 mM of lactose were added. For single template experiments, 0.5 nM of either *ori-Op2Op3-lacZ* (WT) or *ori-Op2Op3-lacZ_E538Q* (MUT) were utilized. In enrichment experiments, a 1:1 DNA mixture (0.25 nM WT and 0.25 nM MUT) or a 1:4 DNA mixture (0.1 nM WT and 0.4 nM MUT) as quantified by Qubit were used.

Liposomes were formed from lipid-coated beads as detailed above. When using single DNA templates (Fig. 4e), 8.5 µL of the liposome suspension were mixed with 1.1 U of DNase I. In the allolactose diffusion experiment (Fig. S10), the *ori-Op2Op3-lacZ* and *ori-Op2Op3-lacZ_E538Q* liposome populations were mixed in a 1:1 ratio (4.3 µL of each) and treated with 1.1 U of DNase I. Reactions were incubated at 30 °C for 16 h (end-point sample), taking a sample after 20 min for start-point measurement. DNase I was then heat inactivated at 75 °C for 15 min and samples were diluted 100-fold in Milli-Q water.

For enrichment experiments, the *ori-Op2Op3-lacZ* and *ori-Op2Op3-lacZ_E538Q* templates were pre-mixed before liposome formation (Fig. 5, Fig. S11). Then, a 10.5-µL liposome solution was treated with 1.3 U of DNase I. After 20 min at 30 °C, a sample was taken (start-point sample) and the incubation was continued for up to 6 h (end-point sample). After DNase I inactivation, samples were diluted 100-fold in Milli-Q water.

Diluted samples were subjected to qPCR analysis or sequenced using long-read Nanopore technology. Prior to sequencing, DNA was PCR amplified using the KOD polymerase. The reaction mix contained 25 µL of Extreme Buffer, 0.02 U µL^−1^ KOD polymerase, 5 µL of diluted liposome sample, 300 nM of primers ChD-980 and 1028, 0.4 mM dNTPs, and 14 µL of Milli-Q water. The thermal cycling program included an initial denaturation step at 94 °C for 2 min, followed by 25 cycles of denaturation at 98 °C for 10 s, annealing at 60 °C for 20 s, and extension at 68 °C for 4 min. The PCR amplicon covering part of *p3* and the complete *lacZ* transcriptional unit was purified using the QIAquick PCR Purification Kit (Qiagen) and sequenced with Nanopore technology. The per-base data file was used to calculate the fraction of reads corresponding to the wild-type or the mutant DNA sequence at position 1612 of the *lacZ* gene. WT: E538 (<u>G</u>AA) and MUT: Q538 (<u>C</u>AA).

With these fraction values, and assuming no replication of the mutated template in the presence of LacI and lactose, we could estimate the overall amplification fold of the DNA mixture using the following equation: *f*_i_MUT_ × (*f*_f_WT_ / *f*_f_MUT_ + 1), where *f*_i_MUT_ refers to the initial fraction of mutated template, *f*_f_WT_ refers to the final fraction of wild-type template, and *f*_f_MUT_ to the final fraction of mutated template. Amplification folds for each starting WT:MUT ratio can be found in Table S1 and be compared with the amplification folds derived from qPCR targeting the *p2* region. In the absence of LacI and lactose, amplification folds cannot be calculated from the fraction values obtained by DNA sequencing.

### IVTTR reactions with gene-encoded Gmk

IVTTR reactions were assembled using PURE*frex*-M1 as described above, except that dGTP in the dNTP mix was replaced with dGMP. The modified dNTP mix was prepared by combining 10 mM each of dATP, dCTP, and dTTP (Thermo Fisher) with 10 mM dGMP (MERCK). A 1.2-µL volume of this dNTP mixture was added per 20-µL of IVTTR reaction, resulting in a final concentration of 600 µM for each deoxynucleotide.

Liposomes were formed as detailed above using lipid-coated beads, either without cholesterol (DOPC, DOPE, DOPG, and cardiolipin in a 50:36:12:2 molar ratio) or with cholesterol (DOPE, DOPG, cardiolipin, and cholesterol in a 35:25:8.5:1.5:30 molar ratio). Cholesterol was purchased from Sigma-Aldrich and dissolved in chloroform. Cholesterol-lipid beads were freshly prepared before use, as we observed that IVTTR activity was compromised when the beads were stored at –20 °C for more than two days. DNA templates consisted of either 0.5 nM of *ori-tag-p2p3* or *ori-gmk-p2p3* for experiments with a single DNA template, or a 1:1 DNA mixture (0.25 nM of each) for competition experiments. In the dGTP diffusion experiment, the *ori-tag-p2p3* and *ori-gmk-p2p3* liposome populations were mixed in a 1:1 ratio. In all cases, 8.5 µL of liposome suspension was treated with 1.1 U of DNase I. After 20 min at 30 °C, a sample was taken (start-point sample) and the incubation was continued for up to 6 h (end-point sample). After DNase I inactivation for 15 min at 30 °C, samples were diluted 100-fold in Milli-Q water prior qPCR analysis.

### Estimating DNA co-encapsulation in liposomes

To assess the probability of multiple DNA molecules being encapsulated within the same liposome, we modeled the DNA encapsulation process using Poisson statistics. The expected number of DNA molecules per liposome, lambda (*λ*), is given by:

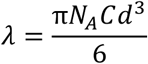

where *N_A_* is the Avogadro’s number, *C* is the bulk input DNA concentration, and *d* is the liposome diameter. The probability of exactly *k* DNA molecules being encapsulated within a liposome according to a Poisson distribution is:

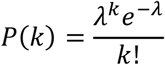

Liposomes prepared using the swelling method exhibit a heterogeneous size distribution, which can be fitted to a log-normal function^6^:

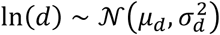

where *μ_d_* and *σ_d_* are the mean and standard deviation of the log-transformed liposome diameter. Experimentally obtained values of *μ_d_* and *σ_d_* are 4.5 µm and 2.3 µm, respectively^6^. Since the encapsulation volume scales with *d*^3^, the expected number of DNA molecules per liposome (*λ*) also follows a log-normal distribution. Therefore, the probability of encapsulating *k* molecules across a population of liposomes is obtained by integrating over the distribution of *d*:

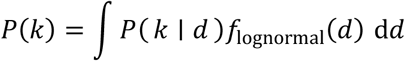

where:

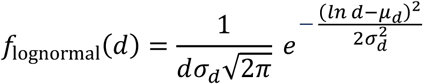

Figure S5 presents the probability of encapsulating 0, 1, 2, 3 or more DNA molecules as a function of DNA concentration considering the measured liposome size variability.

For competition experiments involving two DNA species (e.g., *ori-tag-p2p3* and *ori-T7-p2p3*) at a 1:1 ratio, the probability of at least one copy of each species coexisting within a liposome is given by:

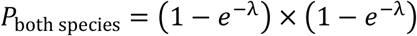

Figure S6 illustrates the probability of both DNA species coexisting within the same liposome as a function of DNA concentration.

### Statistics and reproducibility

All experiments have been replicated independently. The number of replicates is specified in the figure legends. Representative images of gels and colorimetry assays are displayed in the main text figures. Examples of repeated measurements are reported in Supplementary Information.

### Data and code availability

Data are available in the main manuscript and the Supplementary Information. Source data are provided with this paper. Raw data including uncropped gel images are available within a zipped folder named ‘Source Data’. The code used to generate Figs. S5 and S6 and the plasmid maps listed in Table S4 are available on the Github repository (https://github.com/DanelonLab/ACS).

## Acknowledgements

We would like to thank Alicia del Prado and Miguel de Vega (Centro de Biología Molecular Severo Ochoa, Madrid) for kindly providing the purified SSB and DSB proteins, Adilya Timmers (TBI, Toulouse) for plasmid pRS316, Sébastien Nouaille (TBI, Toulouse) for plasmid pBADMyc-HIS-lacZ, Thomas Gosselin-Monplaisir (TBI, Toulouse) for *E. coli* gDNA, and Sara Castaño Cerezo (TBI, Toulouse) for strain CEN.PK2-1C. We are also grateful to Ana María Restrepo Sierra for discussions on DNA replication, and to GeneFrontier for supporting our research and for fruitful discussions. This work was financially supported by Agence Nationale de la Recherche (ANR-22-CPJ2-0091-01).

## Author contributions

L.S.H. and C.D. designed the experiments and wrote the manuscript. L.S.H. performed the experiments and analysed the data. C.D. acquired funding and conceived the project.

