## Supplementary Information for "Autocatalytic selection of gene functions in vitro"

for

**Content:**

Page 2 Table S1. Amplification folds of the DNA mixture for the β-galactosidase competition assay

Page 3 Table S2. List of primers used for PCR

Page 5 Table S3. List of primers used for qPCR

Page 6 Table S4. List of plasmids

Page 7 Figure S1. Optimization of PURE composition for IVTTR reactions

Page 8 Figure S2. Workflow for constructing linear DNA self-replicators with integrated genes using yeast as a cloning platform

Page 9 Figure S3. Biological replicates of protein expression from *ori-tag-p2p3* or *ori-T7-p2p3* DNA template

Page 9 Figure S4. Biological replicates of PURE reactions for assessing activity of gene-encoded T7 RNAP

Page 10 Figure S5. Probability distributions for different numbers of DNA molecules encapsulated within the same liposome at varying DNA concentrations

Page 10 Figure S6. Probability of co-encapsulation of two DNA species in the same liposome

Page 11 Figure S7. Encapsulation and expression of the wild-type *lacZ* transcriptional unit in liposomes at increasing DNA concentrations

Page 12 Figure S8. Biological replicates of protein expression from *ori-Op2Op3-lacZ* (wild-type) or *ori-Op2Op3-lacZ_E538Q* (mutant) DNA template

Page 13 Figure S9. Background β-galactosidase activity in PURE system

Page 13 Figure S10. Assessing diffusion of allolactose between liposomes

Page 14 Figure S11. Autocatalytic selection of an active β-galactosidase encoded in the DNA self-replicator

Page 14 Figure S12. Biological replicates of protein expression from *ori-tag-p2p3* or *ori-gmk-p2p3* DNA template

### Supplementary tables

#### Table S1. Amplification folds of the DNA mixture for the β-galactosidase competition assay

| Ratio (WT:MUT) | + LacI + Lactose | Amplification folds measured by qPCR targeting the *p2* region | Amplification folds estimated from sequencing data |
|---|---|---|---|
| 1:1 | No | 4.30 | N/A |
| 1:1 | Yes | 1.23 | 1.58 |
| 1:4 | No | 5.40 | N/A |
| 1:4 | Yes | 1.35 | 1.35 |

### Table S2. List of primers used for PCR

| Primer pair | DNA sequence (5'→3') | Purpose |
|---|---|---|
| ChD-1365 | TTTGTTTAACTTTAAGAAGGAGATATACATATGAACACGATTAACATCGCTAAGAAC | Amplify the *T7 RNAP* gene from G442. |
| ChD-1366 | AAAAAACTAACAGATACTCTAGTACAATTCTTACGCGAACGCGAAGTCC | |
| ChD-1363 | GAATTGTACTAGAGTATCTGTTAGTTTTTTTCTACTAGAGTACTAGAG | Linearize G435 to clone the *T7 RNAP* gene or the *gmk* gene under PURE regulatory sequences. |
| ChD-1364 | ATGTATATCTCCTTCTTAAAGTTAAACAAAATTATTTCTAGAGG | |
| ChDT-1 | GAATTGTGAGCGGATAACAATTCCCCGGGAGACCACAACGGTTTCC | Linearize G435 with *lacO* overhangs upstream *p2*. |
| ChDT-2 | TTGTTATCCGCTCACAATTCCCCTATAGTGAGTCGTATTAATTTCAC | |
| ChDT-19 | CCCTCTGGAGACACCAGAGG | Linearize pT001 to introduce a *lacO* site upstream *p3*. |
| ChDT-20 | CCCTATAGTGAGTCGTATTAGCAGTCG | |
| ChDT-21 | TACGACTCACTATAGGGGAATTGTGAGCGGATAACAATTCCCCCCCTCTGGAGACACCAG | Generate a dsDNA with a *lacO* site and overhangs to the linearized pT001. |
| ChDT-22 | CTGGTGTCTCCAGAGGGGGGGAATTGTTATCCGCTCACAATTCCCCTATAGTGAGTCGTA | |
| ChDT-7 | TAGCATAACCCCTTGGGGC | Linearize G435 to clone *lacZ* under PURE regulatory sequences. |
| ChDT-8 | TAAACAAAATTATTTCTAGACCCTATAGTGAGTCGTATTAATTTCAC | |
| ChDT-11 | TTTGTTTTGGGTATGACCATGATTACGGATTCACTGGCCGTCGTTTTACAACGTCGTG | Amplify *lacZ* from pBADMyc-HIS-lacZ with start and stop codons. |
| ChDT-12 | CGCCGTAGTCCCAATGAAAATTATTTTTGACACCAGACCAACTGG | |
| ChDT-9 | GAAATAATTTTGTTTAACTTTAAGAAGGAGATATACATATGACCATGATTACGGATTCAC | Add overhangs to the *lacZ* gene amplicon for assembly into the G435 backbone. |
| ChDT-10 | GGCCCCAAGGGGTTATGCTATTATTTTTGACACCAGACCAACTG | |
| ChDT-72 | CCTTTGCCAATACGCCCACGCGATGG | Generate the *lacZ* E538Q (CAA > GAA) mutation in pT003. |
| ChDT-73 | GCGTATTGGCAAAGGATCAGCG | |
| ChDT-199 | TTTGTTTAACTTTAAGAAGGAGATATACATATGGCTCAAGGCACGCTTTATATTG | Amplify the *gmk* gene from *E. coli* genomic DNA. |
| ChDT-200 | AAAAAACTAACAGATACTCTAGTACAATTCTCAGTCTGCCAACAATTTGCTG | |

**Table S2. List of primers used for PCR** (continued)

| | | |
|---|---|---|
| ChDT-41 | TTAGGGAGCACATCCATGCCAATAGCTCGACAAGCGGCGAGAGCCTTGCACCTATGCTATATAAAGTAAGCCCCCACCCTCAC | Amplify the *oriL-T7RNAP* and *oriL-gmk* fragments from the in vitro assembly products for in-yeast assembly of pY002 and pY011, respectively. |
| ChDT-42 | TCAGCGTGTTGTAATGATGCGCCATGAATTAGAATGCGTGATGATGTGCAAAGTGCCGTCGTGAGTCGTATTAGCAGTCGACG | |
| ChDT-43 | GACGGCACTTTGCACATCATCACGCATTCTAATTCATGGCGCATCATTACAACACGCTGATAATACGACTCACTATAGGGAGACC | Amplify the *p2p3-oriR* fragment from G435 for in-yeast assembly of pY002 and pY011. |
| ChDT-44 | GCTACATCTTCCGTACTATGCTGTAGTCTCATGGTCGAGTTCTATTGCTGTTCGGCGGCAAAAGTAGGGTACAGCGACAACATACAC | |
| ChDT-41 | TTAGGGAGCACATCCATGCCAATAGCTCGACAAGCGGCGAGAGCCTTGCACCTATGCTATATAAAGTAAGCCCCCACCCTCAC | Amplify the *oriL-Op2Op3* fragment from pT002 for in-yeast assembly of pY001 and pY008. |
| ChDT-47 | TCAGCGTGTTGTAATGATGCGCCATGAATTAGAATGCGTGATGATGTGCAAAGTGCCGTCGGCAAAAAACCCCTCAAGACC | |
| ChDT-44 | GCTACATCTTCCGTACTATGCTGTAGTCTCATGGTCGAGTTCTATTGCTGTTCGGCGGCAAAAGTAGGGTACAGCGACAACATACAC | Amplify the *lacZ-oriR* fragment (from pT003) and the *lacZ_E538Q-oriR* fragment (from the in vitro assembly reaction) for in-yeast assembly of pY001 and pY008, respectively. |
| ChDT-48 | GACGGCACTTTGCACATCATCACGCATTCTAATTCATGGCGCATCATTACAACACGCTGACGCATGTGAAATTAATACGACTCAC | |
| ChDT-56 | TGCCGCCGAACAGCAATAGAACTCGACCATGAGACTACAGCATAGTACGGAAGATGTAGCCCTGGGTCCTTTTCATCACG | Amplify the *CEN6/ARS4-URA3* fragment from pRS316 for in-yeast assembly of pY001 and pY002. |
| ChDT-57 | ATAGCATAGGTGCAAGGCTCTCGCCGCTTGTCGAGCTATTGGCATGGATGTGCTCCCTAATCTGTGCGGTATTTCACACC | |
| ChDT-65 | TGCCGCCGAACAGCAATAGAACTCGACCATGAGACTACAGCATAGTACGGAAGATGTAGCGGTATCAGCTCACTCAAAG | Amplify the *ColE1-Amp-CEN6/ARS4-URA3* fragment from pRS316 for in-yeast assembly of pY008 and pY011. |
| ChDT-57 | ATAGCATAGGTGCAAGGCTCTCGCCGCTTGTCGAGCTATTGGCATGGATGTGCTCCCTAATCTGTGCGGTATTTCACACC | |
| ChDT-079 | CGTCTGTGTCGCATGTGACCTTAGCATCCCTTCCCTTTGC | Amplify a 170-bp *tag* sequence from pRS316. |
| ChDT-080 | CTATAGTGAGTCGTATTAATTGGTGTGGGTTTAGATGACAAGG | |
| ChDT-077 | TCACATGCGACACAGACGAAGC | Linearize G435 to incorporate a *tag* sequence upstream *p2*. |
| ChDT-078 | AATTAATACGACTCACTATAGGGAGACCAC | |
| ChDT-491E | 5’-PHOS/AAAGTAAGCCCCCACCCTCACATGATACC | Generate 5’-phosphorylated linear templates for IVTTR. |
| ChDT-492E | 5’-PHOS/AAAGTAGGGTACAGCGACAACATACACCATTTCC | |
| ChDT-016 | AAAGTAAGCCCCCACCCTCACATG | Amplify the *lacZ* transcriptional unit from pT003. |
| ChDT-017 | AAAGTAGGGTACAGCGACAACATACAC | |
| ChD-980 | ACGGCTGAAATTGACATCCCG | Amplify a region spanning part of *p3* and the *lacZ* transcriptional unit for Nanopore sequencing of diluted IVTTR samples. |
| ChD-1028 | AGGTAATAAGACAACCAATCATAGGAGGCA | |

**Table S3. List of primers used for qPCR**

| Primer pair | DNA sequence (5'→3') | Purpose |
|---|---|---|
| ChD-976<br>ChD-977 | GGATGAAGACTACCCGCTGC<br>ACAGGTCTGCGATTTCACCG | qPCR amplicon in *p2* gene |
| ChDT-81<br>ChDT-82 | CCTTAGCATCCCTTCCCTTTGC<br>GGTGTGGGTTTAGATGACAAGG | qPCR amplicon in *tag* |
| ChDT-52<br>ChDT-53 | CTGGAGAGATTCTTCGCAAGCG<br>GTAAGCGGAACTGACCGAGG | qPCR amplicon in *T7 RNAP* gene |
| ChDT-223<br>ChDT-224 | GCAAATTCGCCAGAAGATGC<br>CTTGCGCCATACGCTTTGC | qPCR amplicon in *gmk* gene |

**Table S4. List of plasmids**

| Plasmid name | Relevant characteristics |
|---|---|
| G435 | *ori-p2p3; AmpR; ColE1*<br>*p2* transcriptional unit: *T7p g10L RBS p2 vsv-r1 vsv-r2*<br>*p3* transcriptional unit: *T7p mod-g10L RBS p3 T7t* |
| G442 | *T7 RNAP* gene; *KanR; oriV* |
| pRS316 | *CEN6/ARS4-URA3; AmpR; ColE1* |
| pBADMyc-HIS-lacZ | *lacZ* gene without start and stop codon; *AmpR; ColE1* |
| pT001 | *ori-Op2p3; AmpR; ColE1*<br>*p2* transcriptional unit: *T7p lacO g10L RBS p2 vsv-r1 vsv-r2* |
| pT002 | *ori-Op2Op3; AmpR; ColE1*<br>*p3* transcriptional unit: *T7p lacO mod-g10L RBS p3 T7t* |
| pT003 | *ori-lacZ; AmpR; ColE1*<br>*lacZ* transcriptional unit: *T7p g10L RBS lacZ T7t* |
| pT007 | *ori-tag-p2p3*; *AmpR; ColE1* |
| **In-yeast assembled plasmids** | |
| pY001 | *ori-Op2Op3-lacZ*; *CEN6/ARS4-URA3* |
| pY002 | *ori-T7-p2p3*; *CEN6/ARS4-URA3*<br>*T7 RNAP* transcriptional unit: *T7p g10L RBS T7 RNAP vsv-r1 vsv-r2* |
| pY008 | *ori-Op2Op3-lacZ_E538Q*; *CEN6/ARS4-URA3; AmpR; ColE1* |
| pY011 | *ori-gmk-p2p3*; *CEN6/ARS4-URA3; AmpR; ColE1*<br>*gmk* transcriptional unit: *T7p g10L RBS gmk vsv-r1 vsv-r2* |

### Supplementary Figures

a)

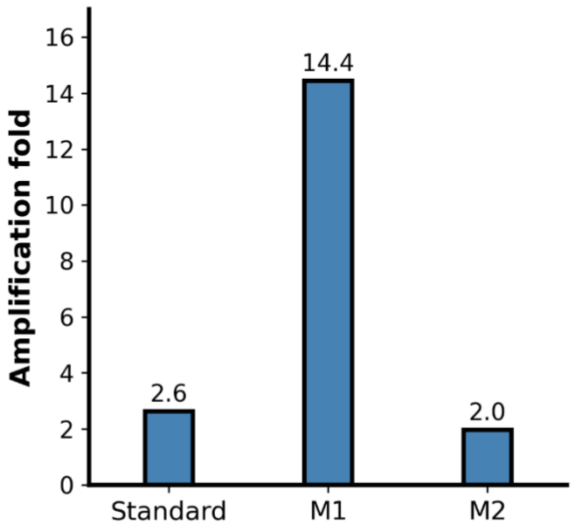


b)

| Reagent name | M1 Factor Modified | M2 Factor Modified |
|---|---|---|
| HEPES | 2.5 | 2.5 |
| K-Glutamate | 0.39 | 0.39 |
| **Mg-acetate** | **0.58** | **1** |
| DTT | 2 | 2 |
| Spermidine | 0.5 | 0.5 |
| 10-fTHF | 1 | 1 |
| CP | 0.42 | 0.42 |
| 18 Aas | 0.6 | 0.6 |
| Tyr | 1 | 1 |
| Cys | 1 | 1 |
| **ATP/GTP** | **0.33** | **1** |
| **CTP/UTP** | **0.33** | **1** |
| tRNA mix | 0.33 | 0.33 |

**Fig. S1. Optimization of PURE composition for IVTTR reactions**. **a**) Amplification folds from bulk IVTTR reactions using the standard PURE*frex*2.0 and two modified Solution I compositions. *Ori-p2p3* at 500 pM was used as template. **b**) Relative changes in concentration of the two modified compositions, M1 and M2, with respect to the standard PURE*frex*2.0. For example, a factor of 2 means that the concentration has been doubled, while a factor of 0.5 means the concentration has been decreased by half. Increases are shown in green and decreases in orange. The difference between M1 (active for IVTTR) and M2 (inactive for IVTTR) lies in the concentration of Mg-acetate and NTPs (bold), indicating these are critical factors for successful replication. Specific working concentrations are not disclosed for confidentiality reasons as agreed with GeneFrontier Corp. The optimized PURE*frex*-M1 is available from GeneFrontier as a customized kit.

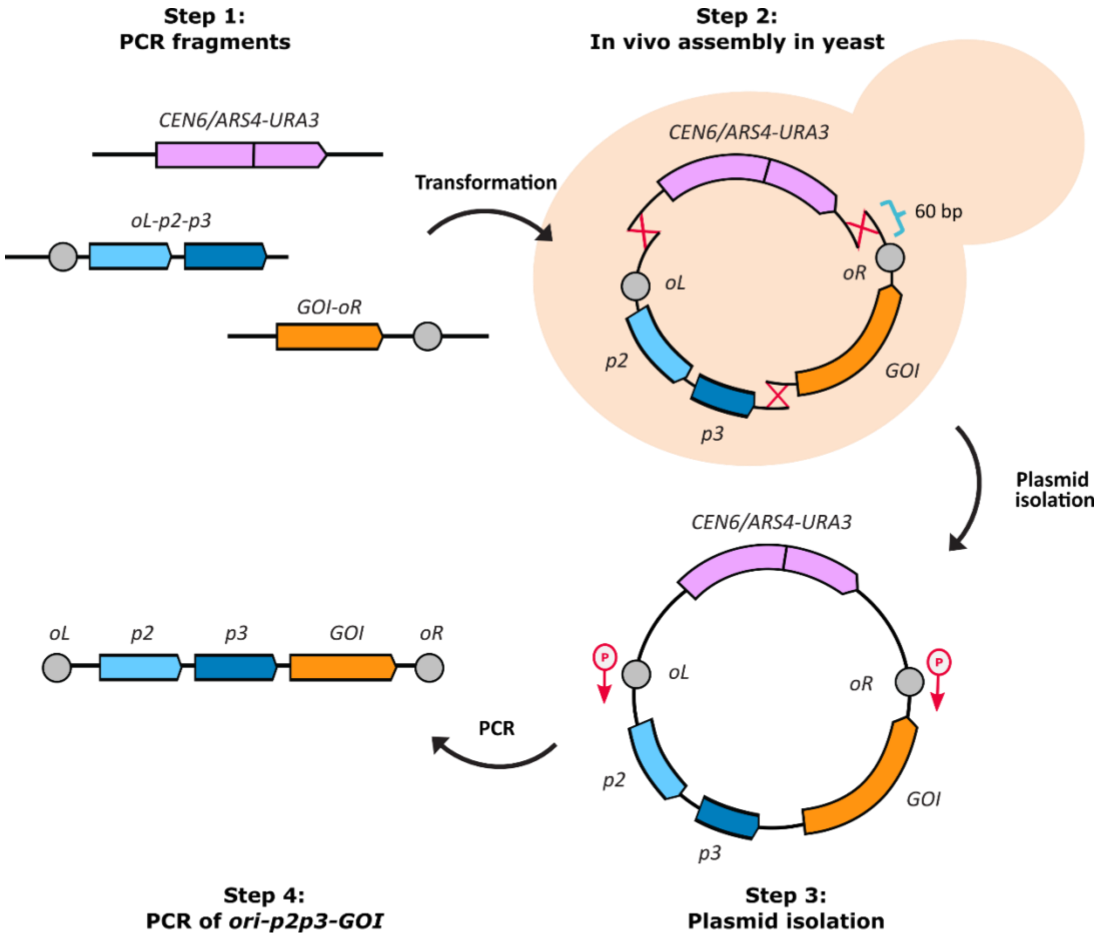


**Fig. S2. Workflow for constructing linear DNA self-replicators with integrated genes using yeast as a cloning platform.** The process consists of four steps: (1) Amplification of the genetic elements via PCR. This includes the *p2-p3* fragment and the gene of interest (GOI) fragment. Pre-cloning the GOI between ϕ29 replication origins ensures that one origin is already part of the fragment. The GOI can be introduced either upstream or downstream of the replication genes. A fragment containing a yeast origin of replication and selection marker (*CEN6/ARS4-URA)* is required for amplification and maintenance in yeast. (2) Transformation of yeast with the DNA fragments for in vivo assembly. The DNA fragments were designed with 60-bp synthetic overlaps for homologous recombination. (3) Isolation of the assembled plasmid. (4) Generation of linear self-replicators through PCR using 5′ phosphorylated primers.

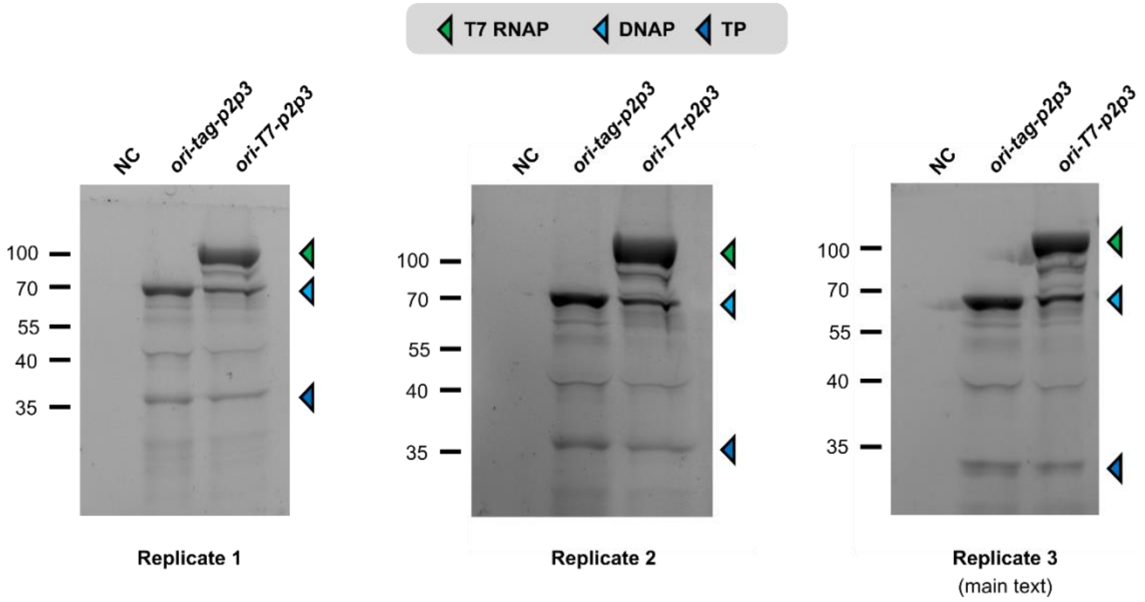


**Fig. S3. Biological replicates of protein expression from *ori-tag-p2p3* or *ori-T7-p2p3* DNA template.** PURE reactions were supplemented with GreenLys reagent and visualized on SDS-PAGE gels. T7 RNAP (~ 99 kDa), DNAP (~ 66 kDa), and TP (~ 31 kDa). NC: negative control with no DNA.

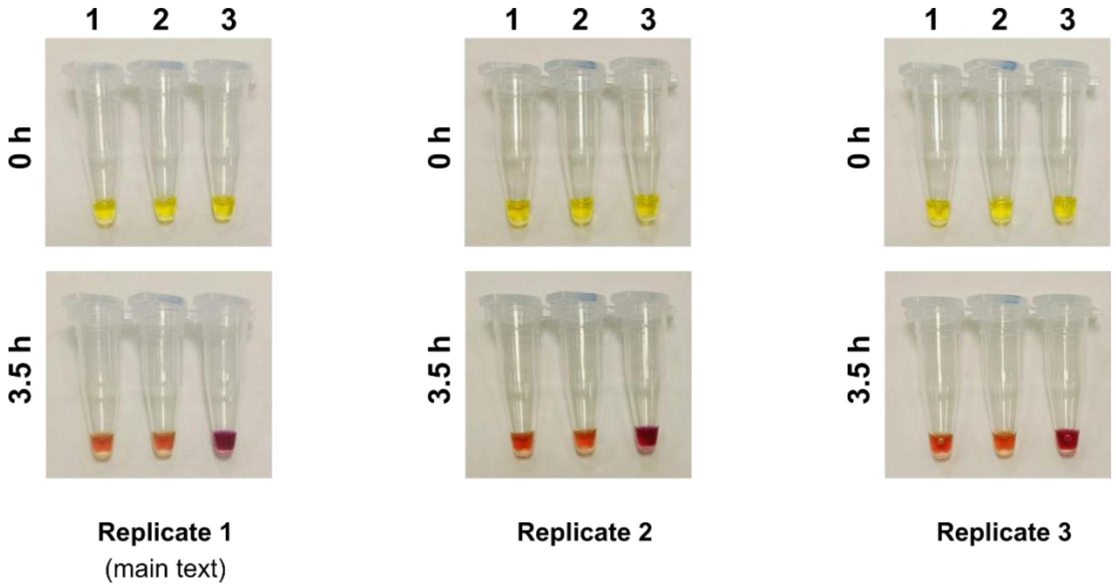


**Fig. S4. Biological replicates of PURE reactions for assessing activity of gene-encoded T7 RNAP.** The experiment was performed as described in Fig. 2c. Images were taken following CPRG addition (0 h) and 3.5 h later. (1) No DNA, (2) *ori-tag-p2p3*, and (3) *ori-T7-p2p3*.

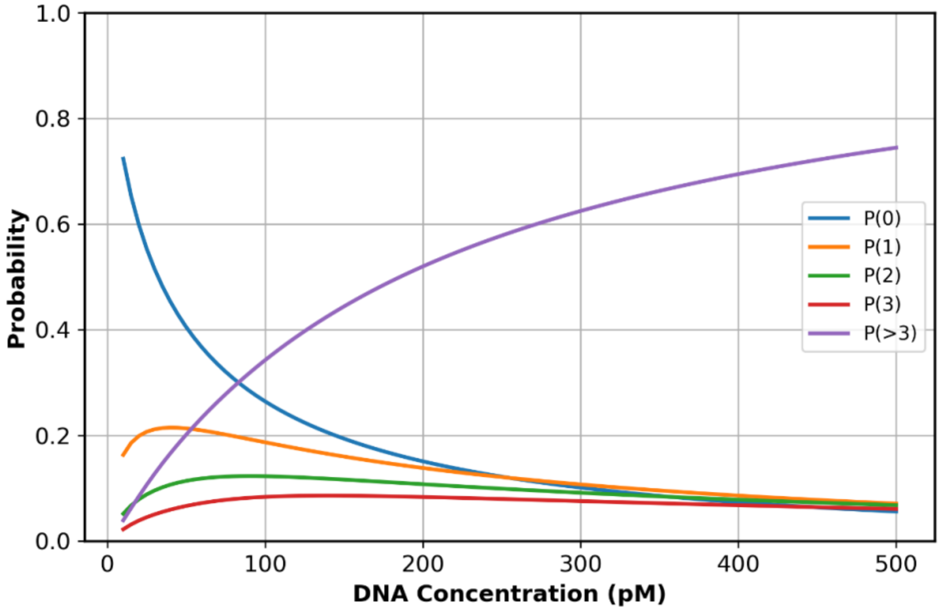


**Fig. S5. Probability distributions for different numbers of DNA molecules encapsulated within the same liposome at varying DNA concentrations.** The probabilities were calculated for 0, 1, 2, 3, and more than 3 molecules using a Poisson distribution. The liposome diameter is modeled using a log-normal distribution with a mean of $\mu_d$ = 4.5 µm and a standard deviation $\sigma_d$ = 2.3 µm.

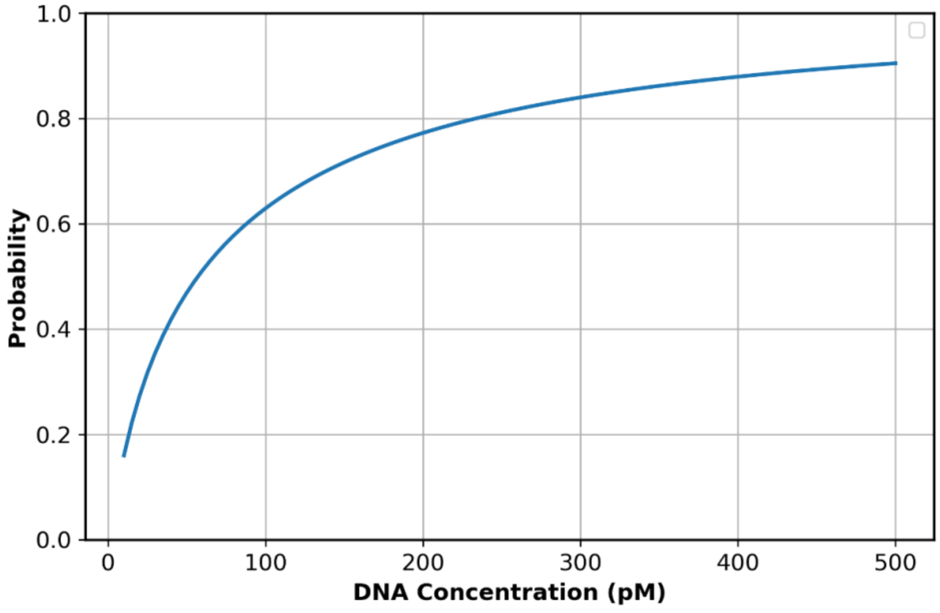


**Fig. S6. Probability of co-encapsulation of two DNA species in the same liposome.** The plot shows the probability of at least one copy of each DNA species being encapsulated within the same liposome, calculated for a 1:1 DNA ratio.

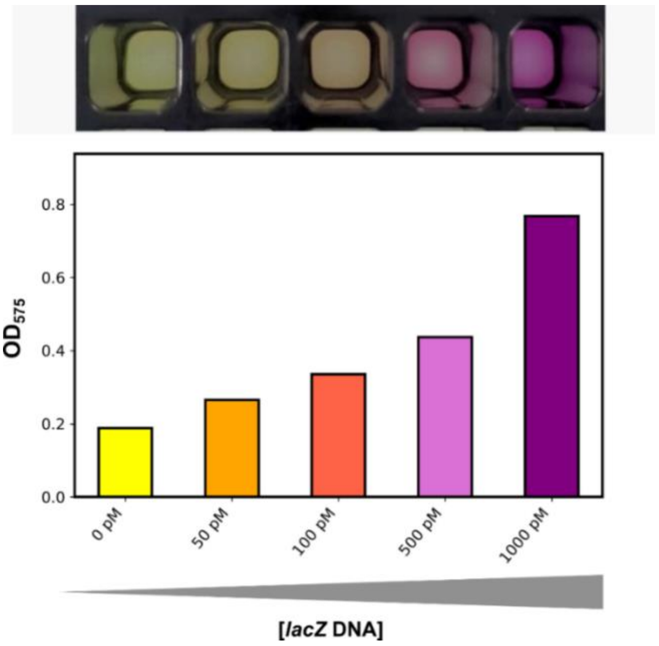


**Fig. S7. Encapsulation and expression of the wild-type *lacZ* transcriptional unit in liposomes at increasing DNA concentrations.** After liposome formation, CPRG was added to monitor β-galactosidase activity. Reactions were incubated for 2 h at 30 °C, after which images were taken (top) and $OD_{575}$ values were measured (bottom). Both the color change and absorbance values correlate with the increasing amount of input DNA.

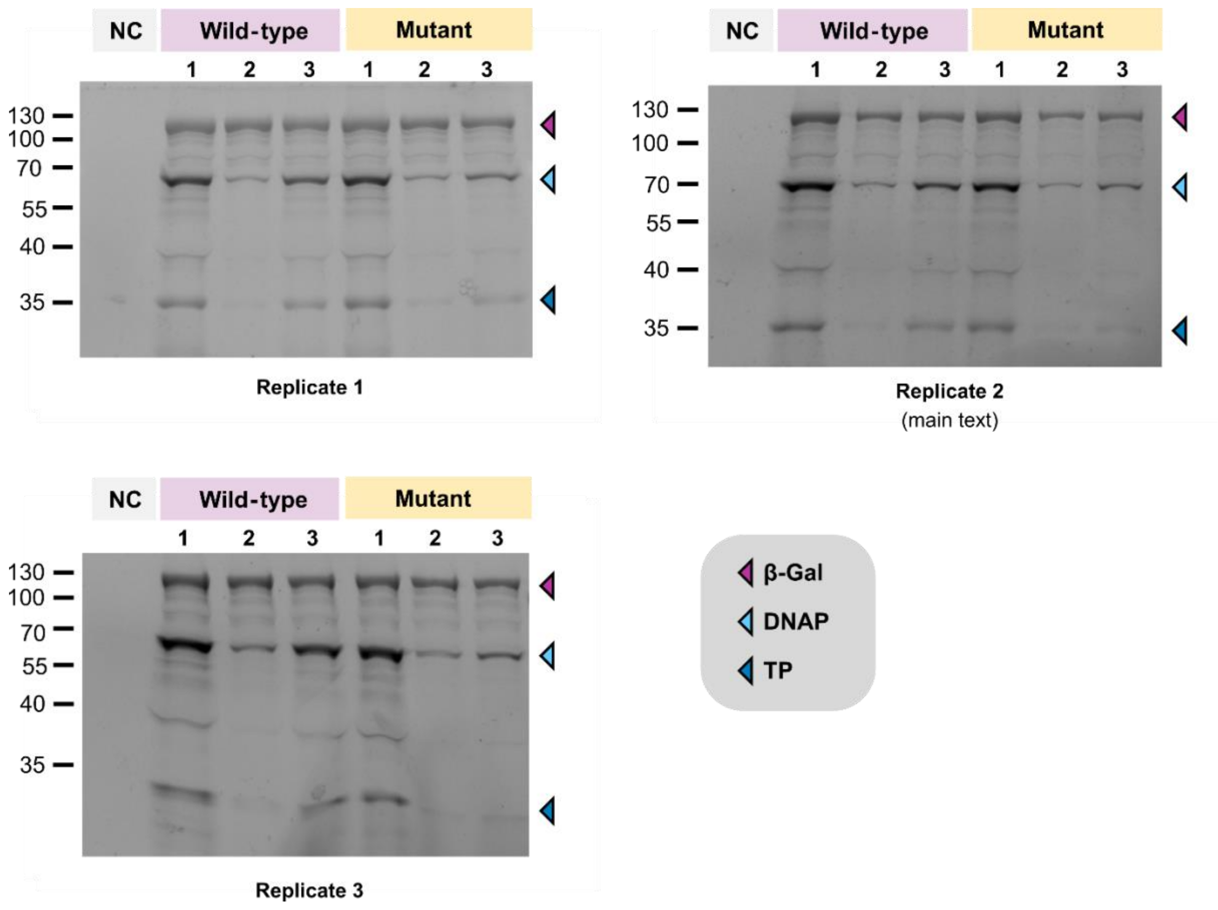


**Fig. S8. Biological replicates of protein expression from *ori-Op2Op3-lacZ* (wild-type) or *ori-Op2Op3-lacZ_E538Q* (mutant) DNA template.** PURE reactions were supplemented with GreenLys reagent and visualized on SDS-PAGE gels. Conditions: (1) in the absence of LacI; (2) in the presence of LacI; (3) in the presence of LacI and lactose. β-galactosidase (~ 117 kDa), DNAP (~ 66 kDa), and TP (~ 31 kDa). NC: negative control with no DNA.

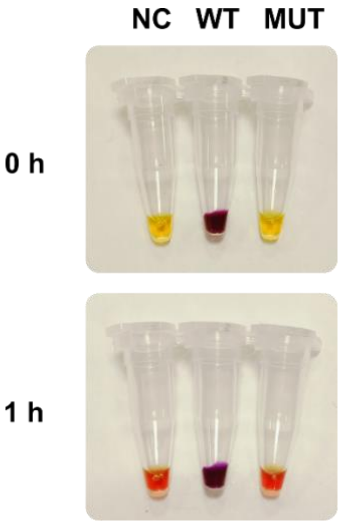


**Fig. S9. Background β-galactosidase activity in PURE system.** A progressive color change is observed in the negative control (NC) sample without a DNA template. The top image corresponds to Fig. 4b (main text) and was taken after 1 h of expression at 30 ˚C and immediately following CPRG addition. The bottom image was taken 1 h later, showing conversion in the NC sample, which indicates the presence of a β-galactosidase-like side reaction in PURE system. Notably, the NC and the mutated (MUT) template samples exhibit the same color change, suggesting that the observed effect is likely due to residual β-galactosidase activity in PURE rather than activity of the mutant enzyme.

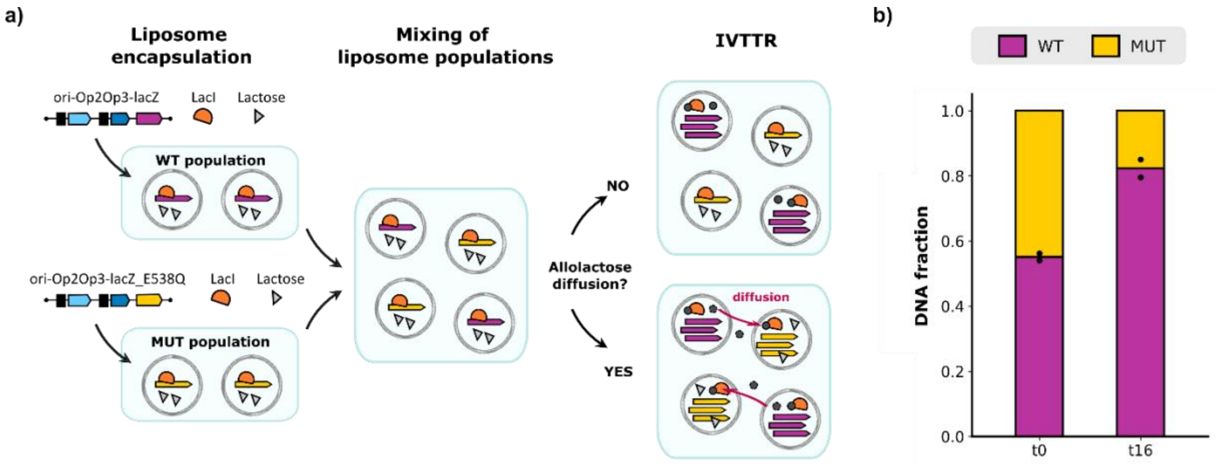


**Fig. S10. Assessing diffusion of allolactose between liposomes. a**) Schematic illustration of the experimental setup to evaluate allolactose diffusion. Two liposome populations were generated: one encapsulating the wild-type DNA template (*ori-Op2Op3-lacZ*) and the other containing the mutated DNA template (*ori-Op2Op3-lacZ_E538Q*), both in the presence of LacI and lactose. The populations were mixed in a 1:1 ratio and incubated for 16 h. In the absence of allolactose diffusion (top), only the wild-type template would replicate, maintaining the genotype-to-phenotype linkage. In contrast, if allolactose diffuses between liposomes, both templates would replicate, compromising the selection process. **b**) Fraction of wild-type (WT) and mutated (MUT) DNA templates in the population before (t0) and after (t16) IVTTR, as determined by long-read sequencing. Enrichment of WT over MUT after IVTTR indicates that allolactose diffusion across the liposome membrane is too low to compromise selection. Individual symbols represent data from two biological replicates (independent experiments).

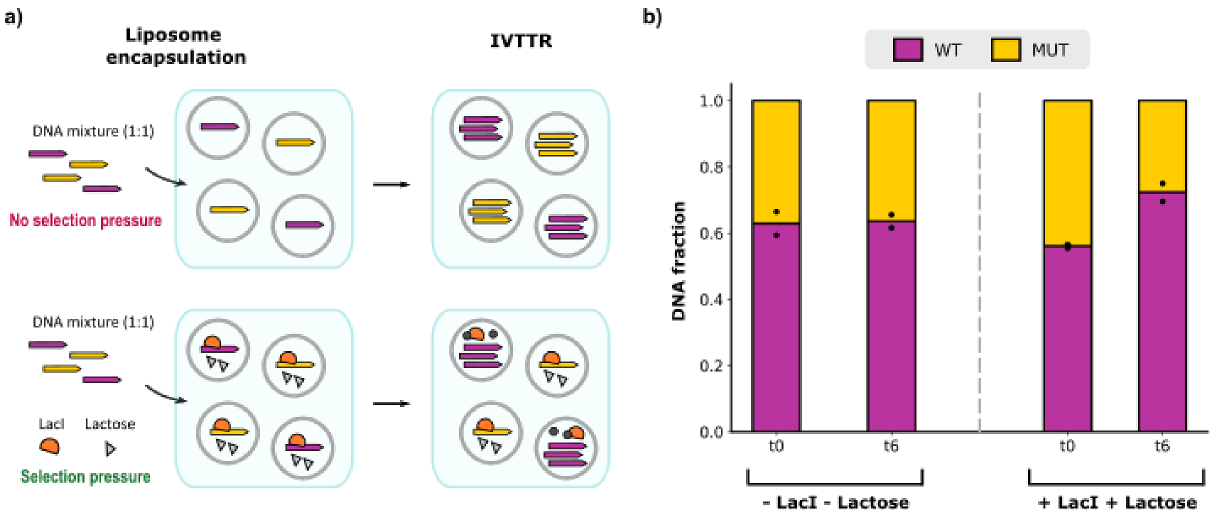


**Fig. S11. Autocatalytic selection of an active β-galactosidase encoded in the DNA self-replicator. a)** Schematic illustration of the experimental conditions and predicted replication outcomes. A 1:1 mixture of wild-type (purple) and mutated (yellow) DNA templates was encapsulated along with the IVTTR components into liposomes (total DNA concentration was 500 pM). In the absence of LacI (no selection pressure), both templates are expected to replicate. When LacI and lactose are co-encapsulated (selection pressure), only the wild-type template is expected to be de-repressed providing a replication advantage. **b**) Fraction of the wild-type (WT) and mutated (MUT) variants in the DNA mixture before (t0) and after (t6) IVTTR. The number of reads for each template, as determined by long-read Nanopore sequencing, was used to quantify the relative abundance of each variant. Individual symbols represent data from two biological replicates (independent experiments).

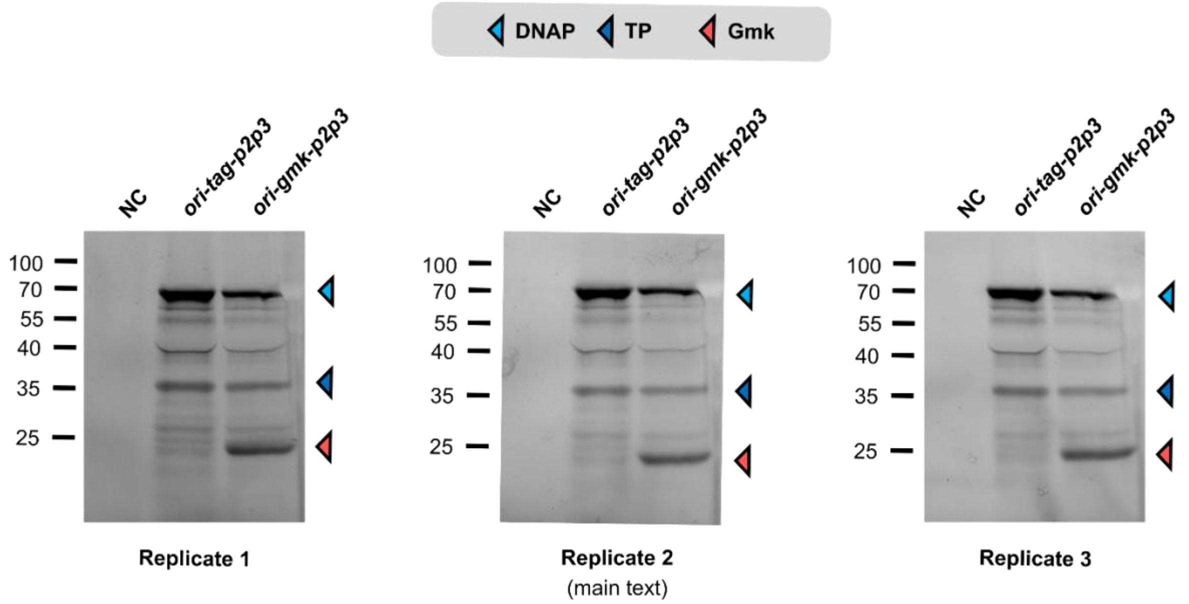


**Fig. S12. Biological replicates of protein expression from *ori-tag-p2p3* or *ori-gmk-p2p3* DNA template.** PURE reactions were supplemented with GreenLys reagent and visualized on SDS-PAGE gels. DNAP (~ 66 kDa), TP (~ 31 kDa), and Gmk (~ 24 kDa). NC: negative control with no DNA.
