## Supplementary figures and images for "Autocatalytic selection of gene functions in vitro"

### 231120_Picture_at_1739.jpeg

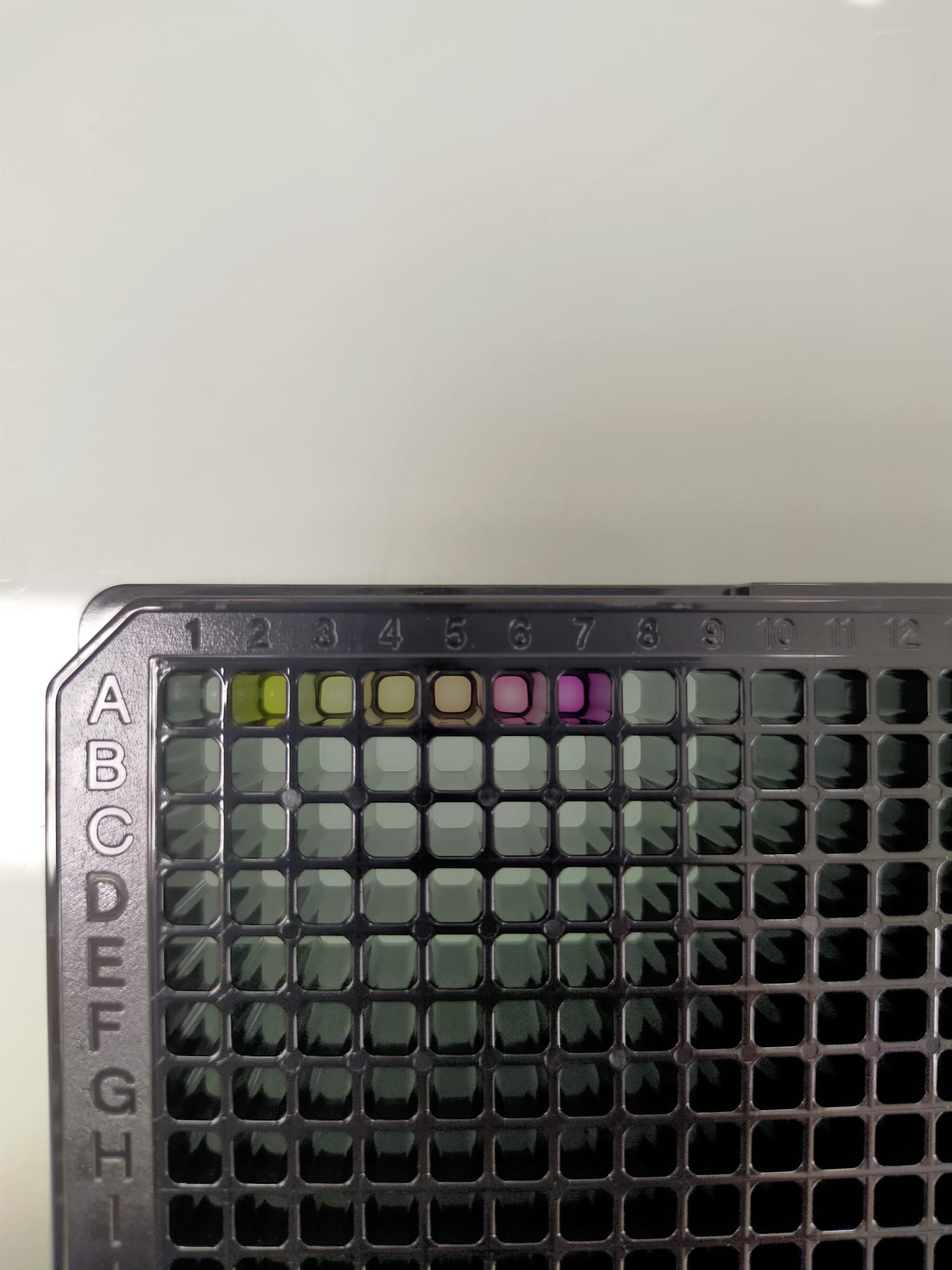

### IMG20241121144610 - CPRG.jpg

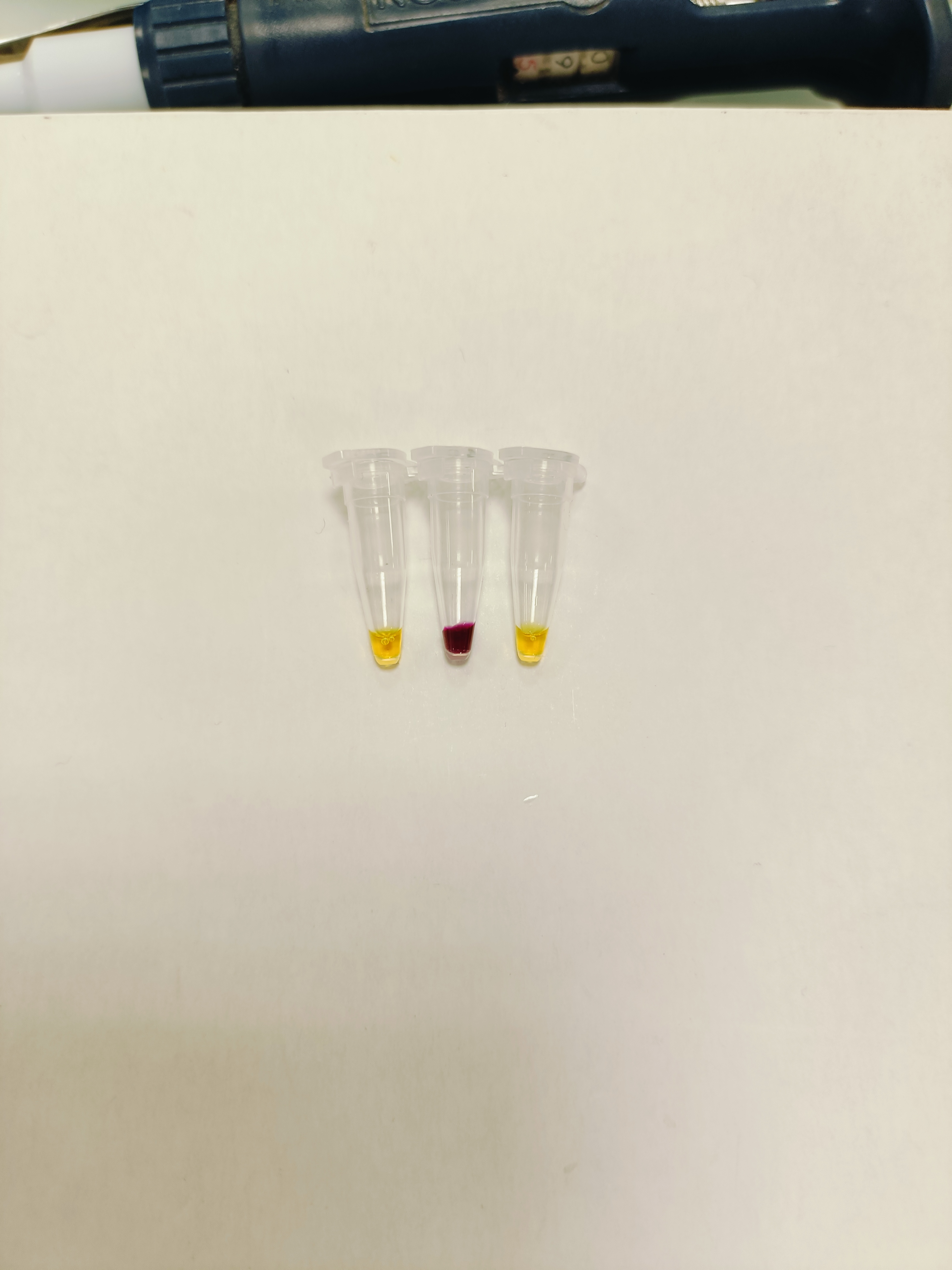

### IMG20241121151337 - Xgal.jpg

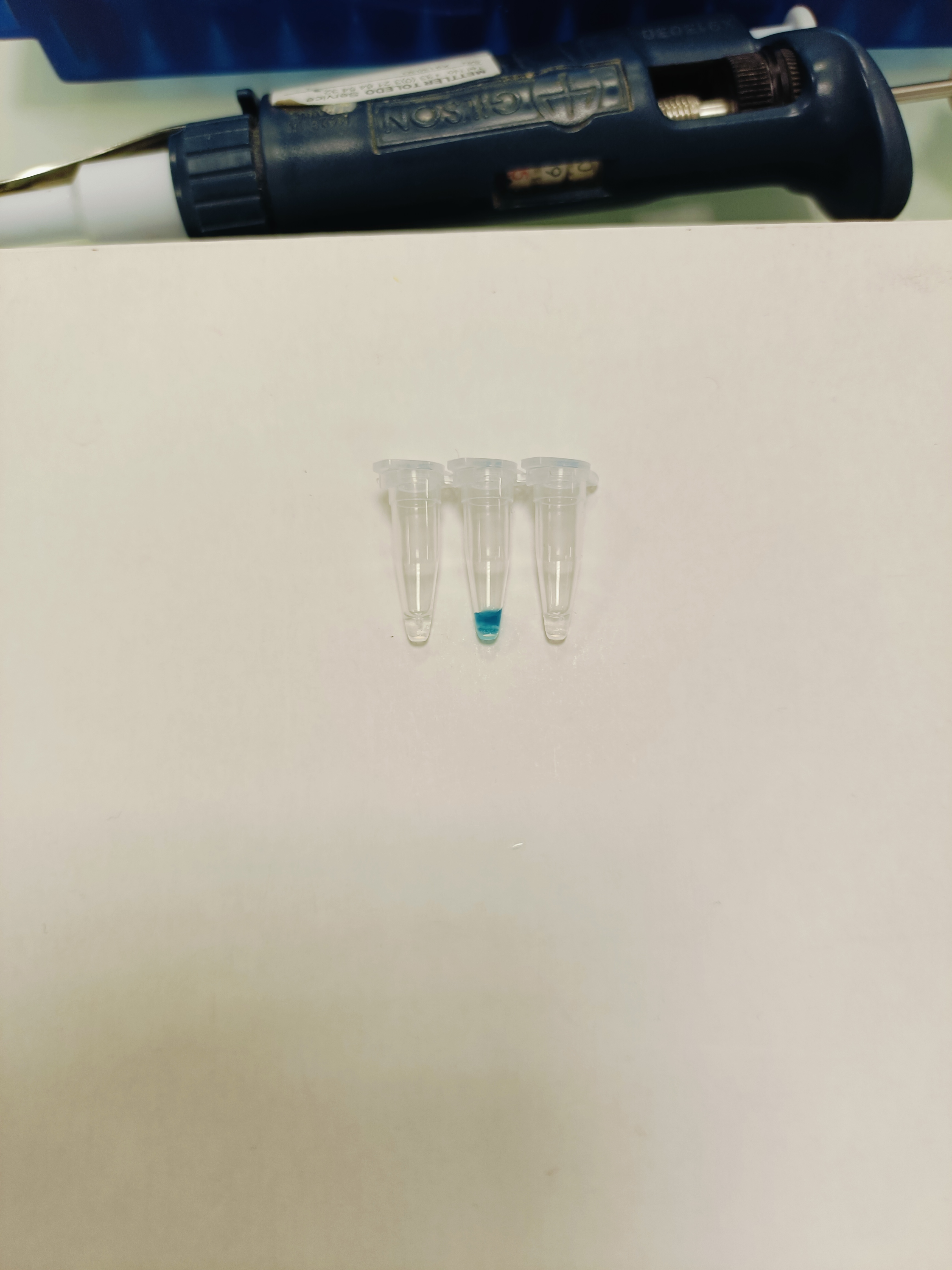

### IMG20241121154132.jpg

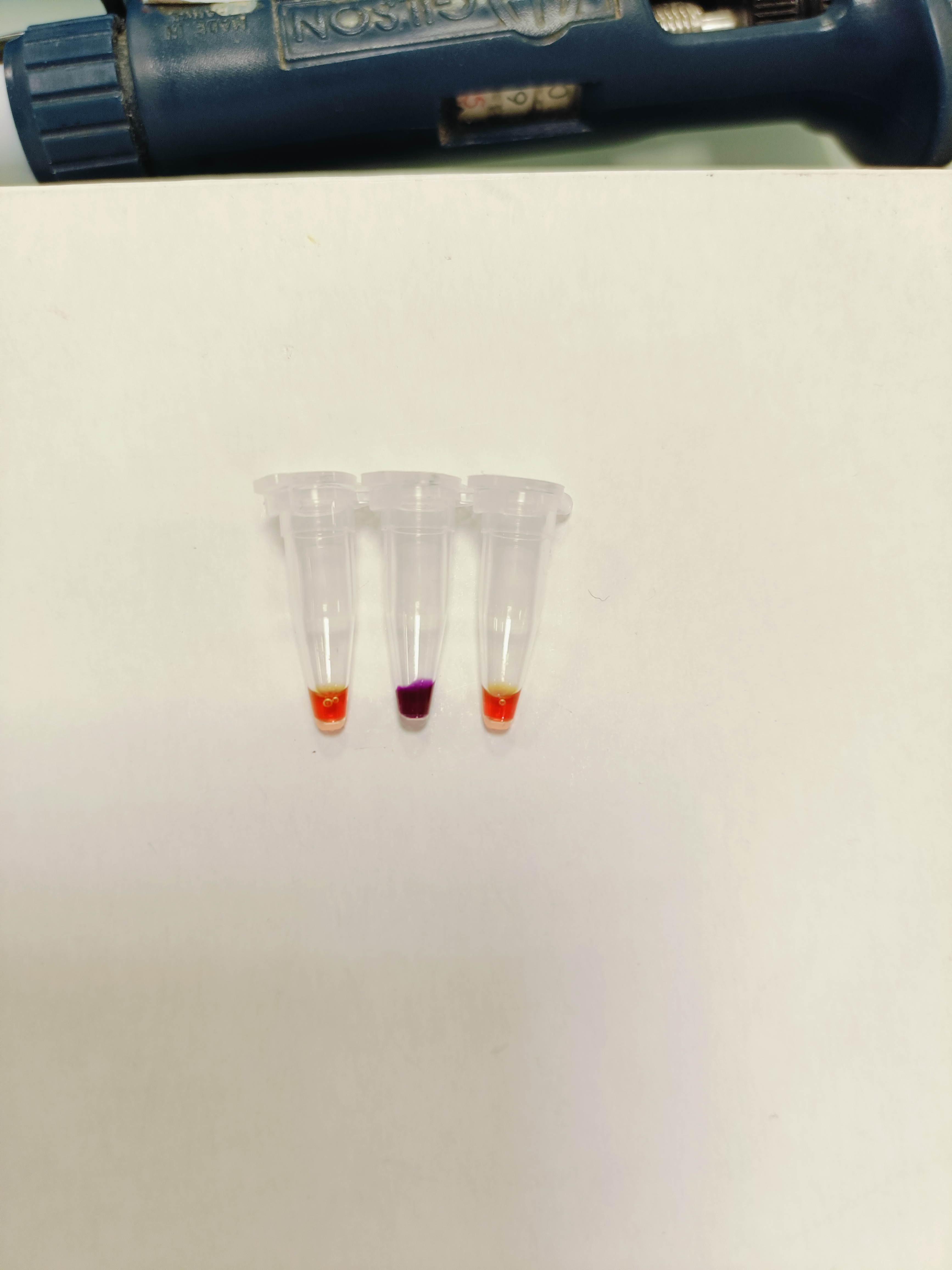

### IMG20241211090624.jpg

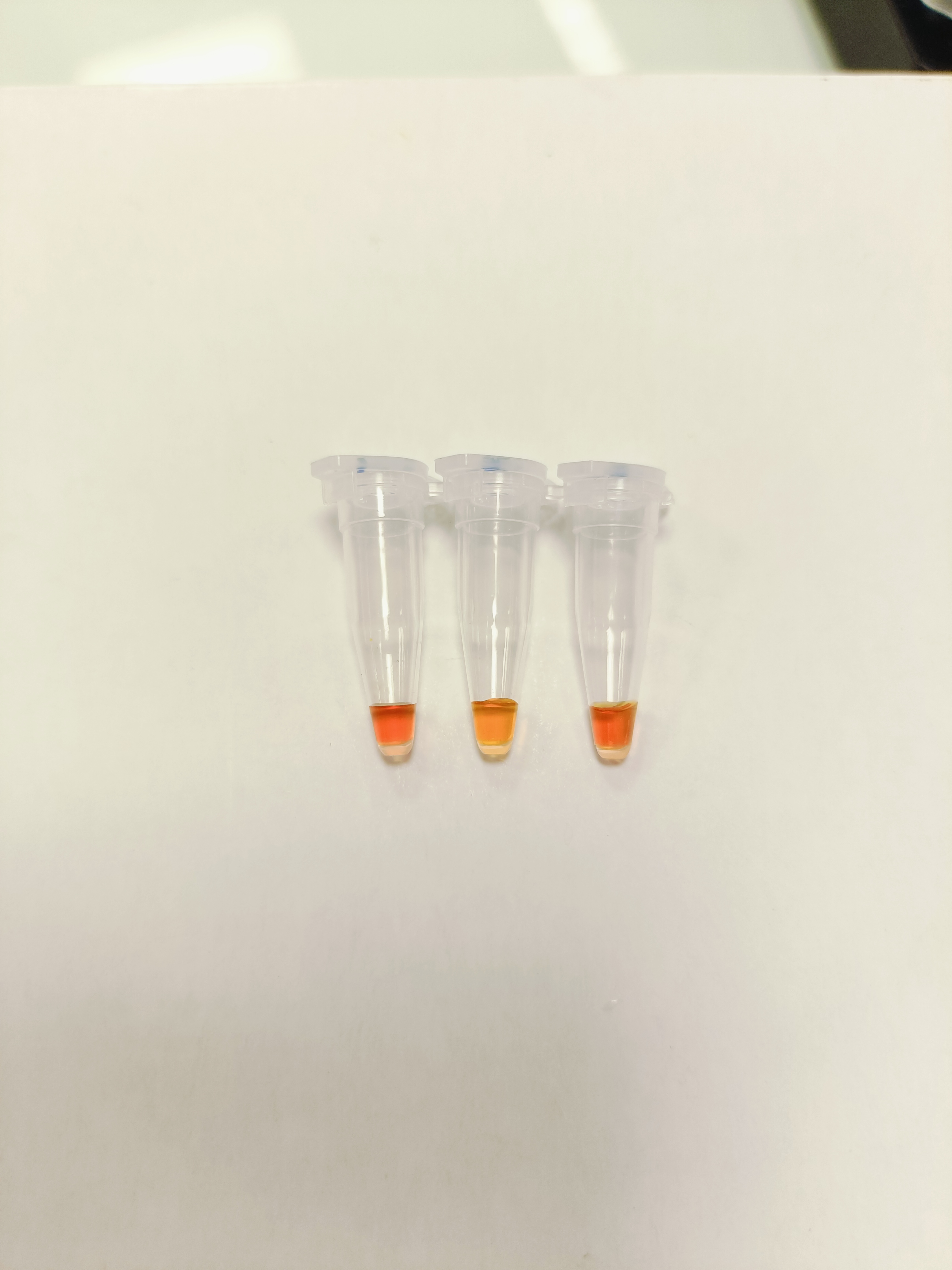

### IMG20241211090633.jpg

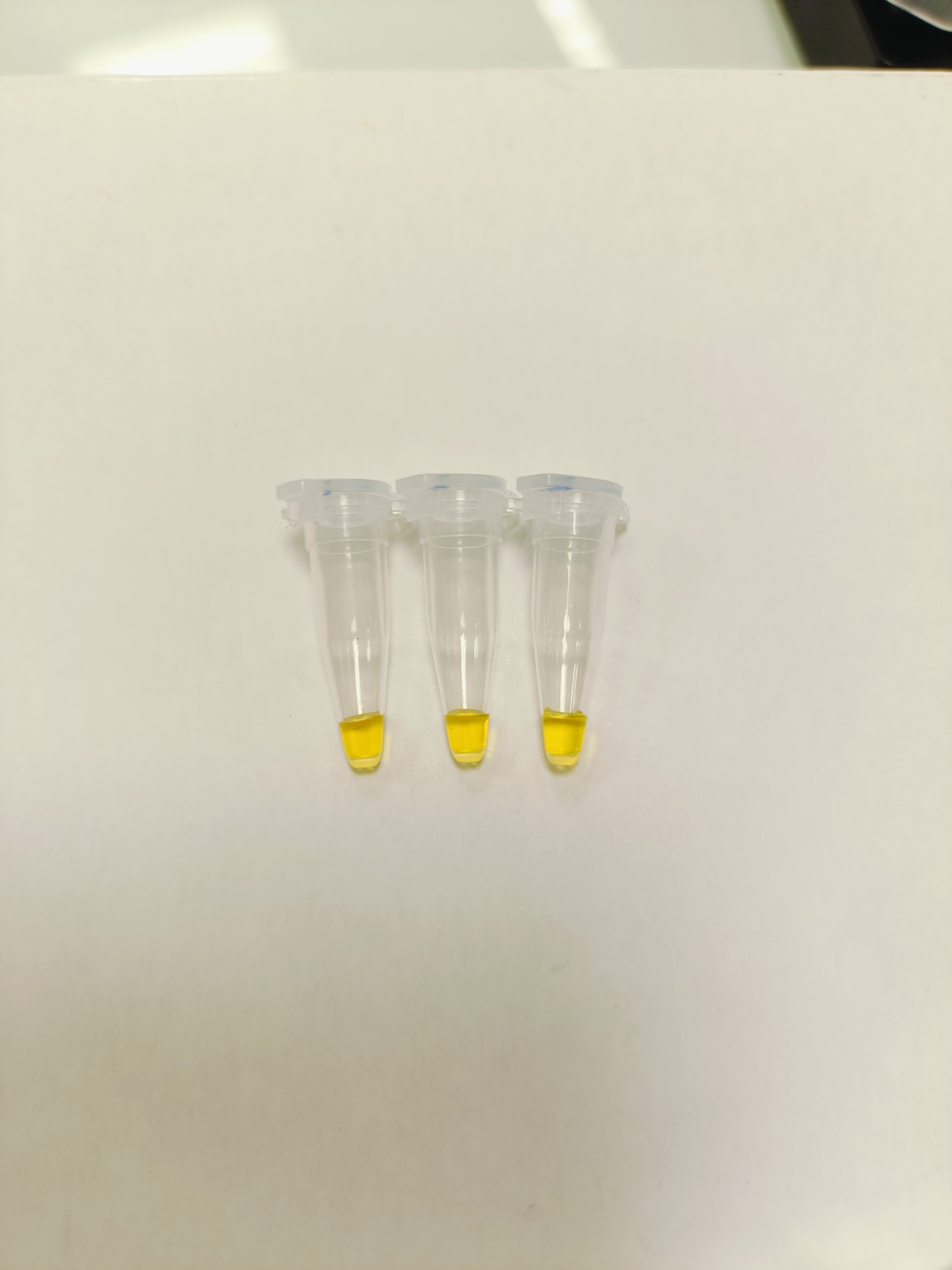

### Rep1 - ead11 - Laura Sierra 2024-09-24 11h21m02s(Alexa 488).tif

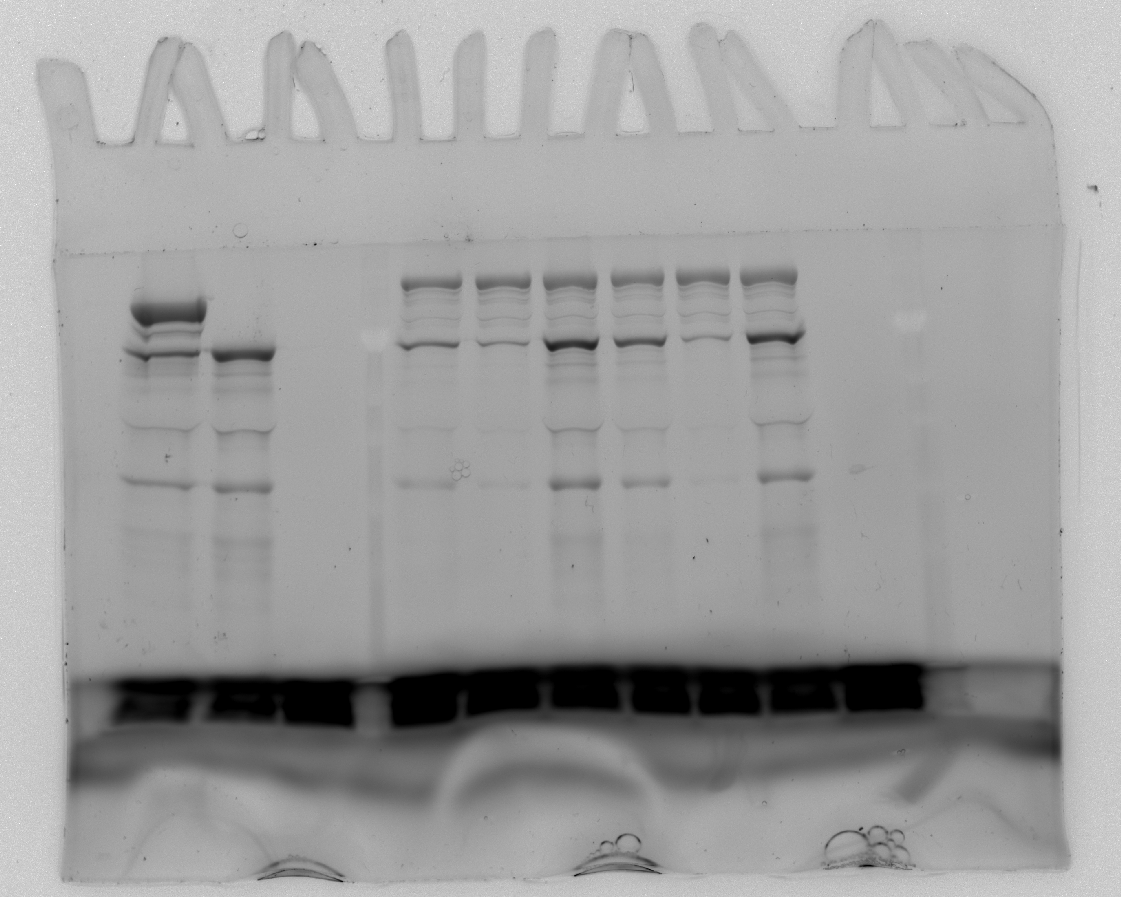

### Rep1 - ead11 - Laura Sierra 2024-09-24 11h21m02s(Composite).tif

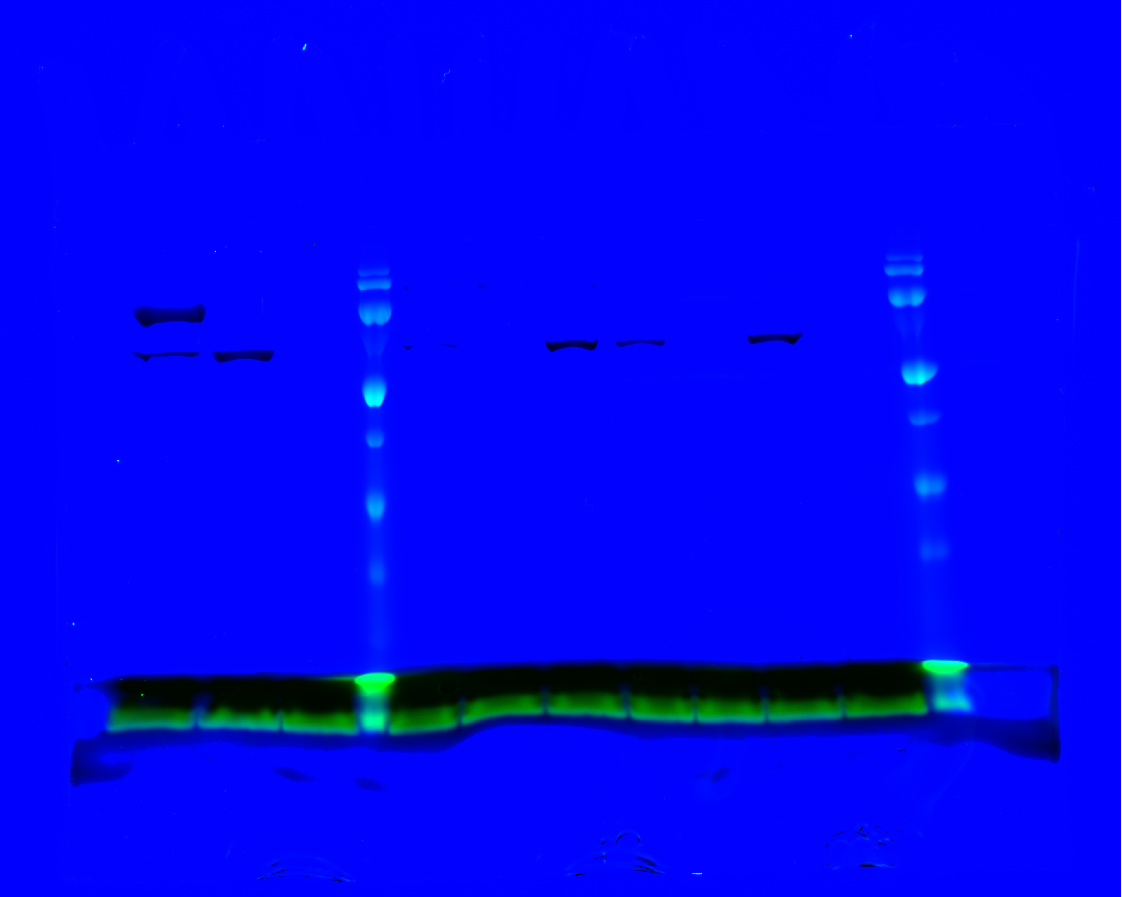

### Rep1 - ead11 - Laura Sierra 2024-12-06 14h48m46s(Alexa 488).tif

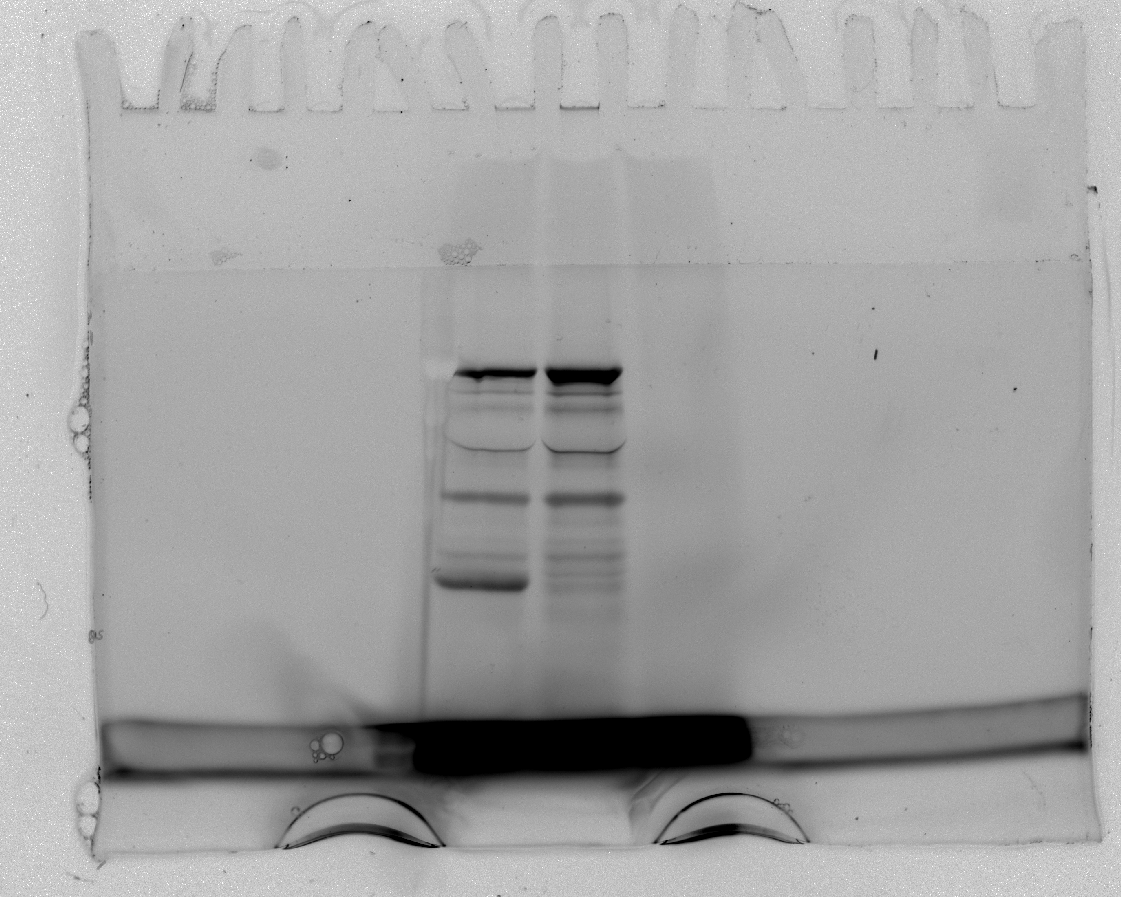

### Rep1_IMG20241011115147.jpg

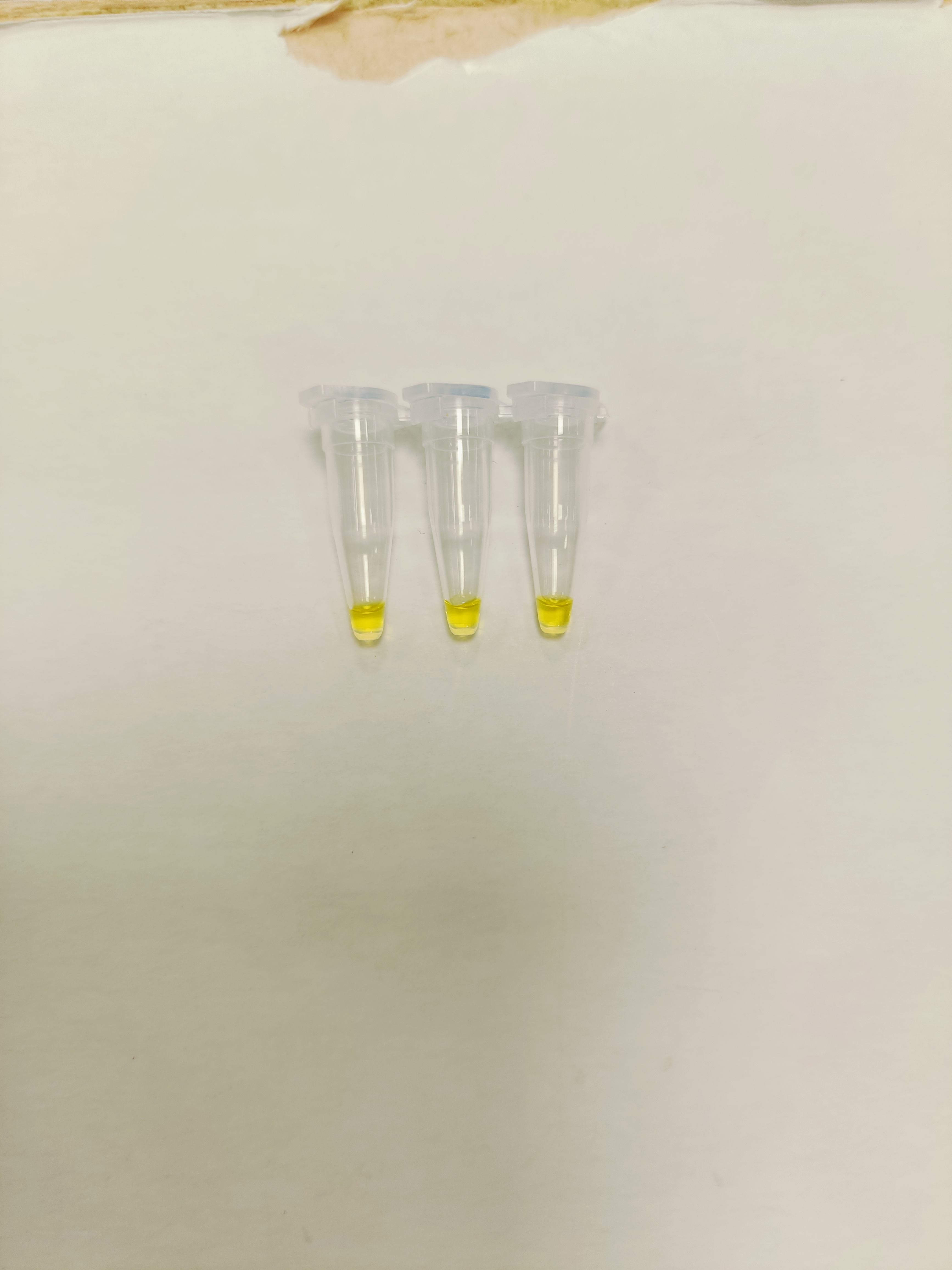

### Rep1_IMG20241011152308.jpg

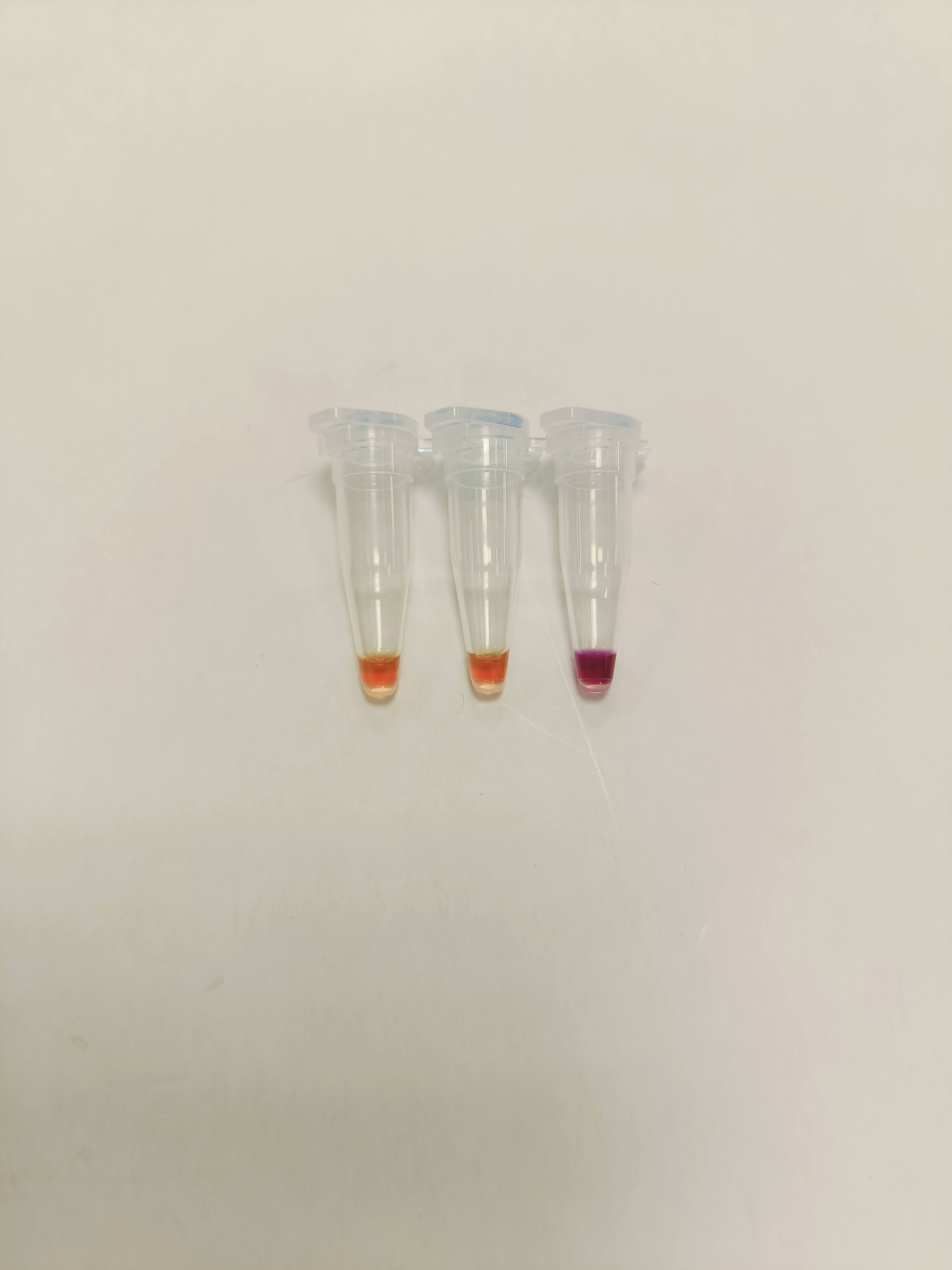

### Rep2 - ead11 - Laura Sierra 2024-10-02 13h34m47s(Alexa 488).tif

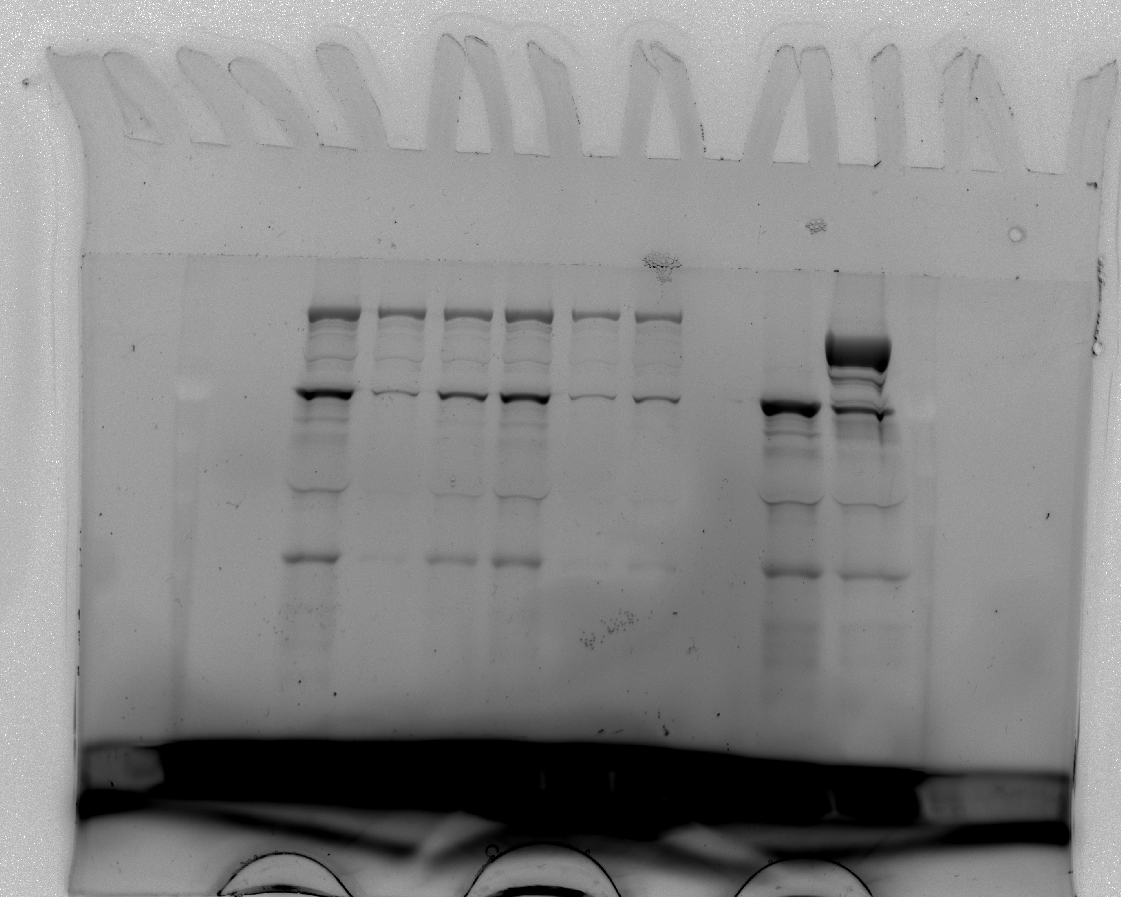

### Rep2 - ead11 - Laura Sierra 2024-10-02 13h34m47s(Composite).tif

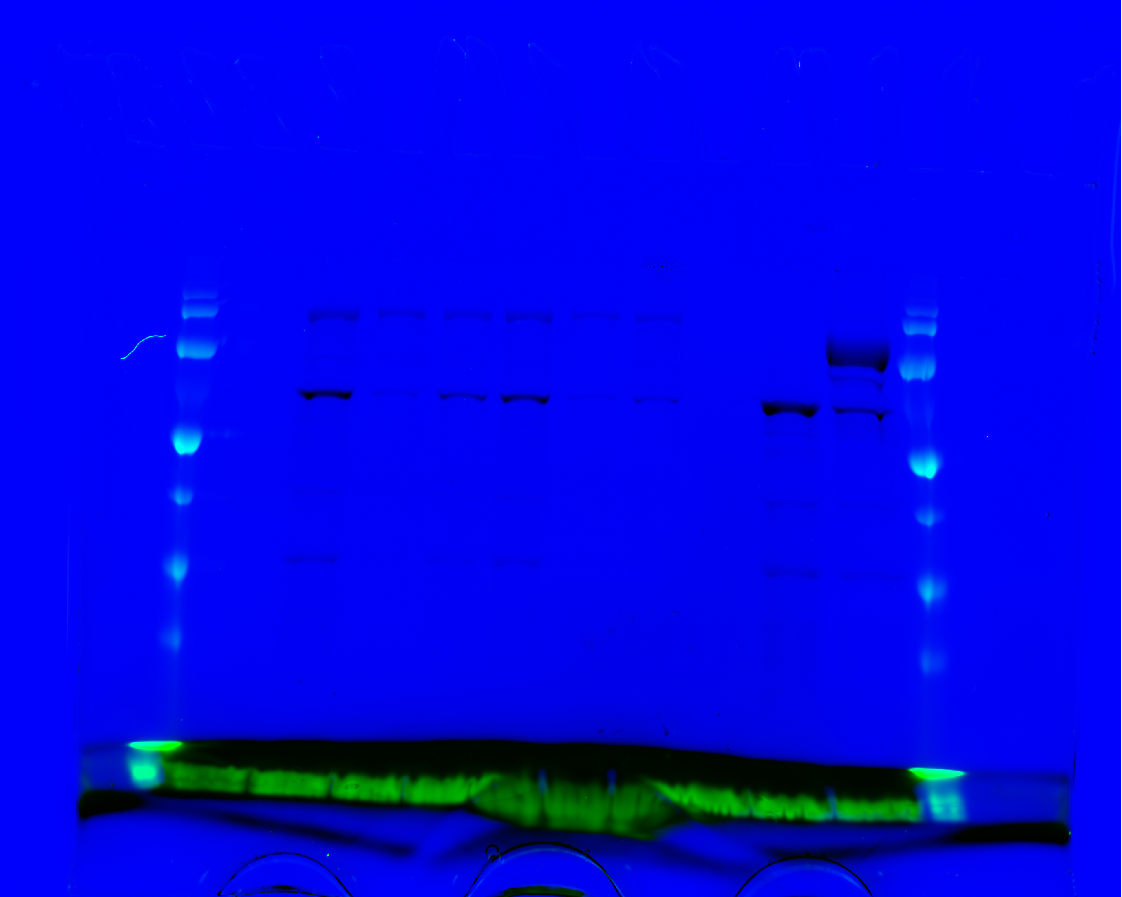

### Rep2 and Rep3 - ead11 - Laura Sierra 2024-12-06 14h56m31s(Alexa 488).tif

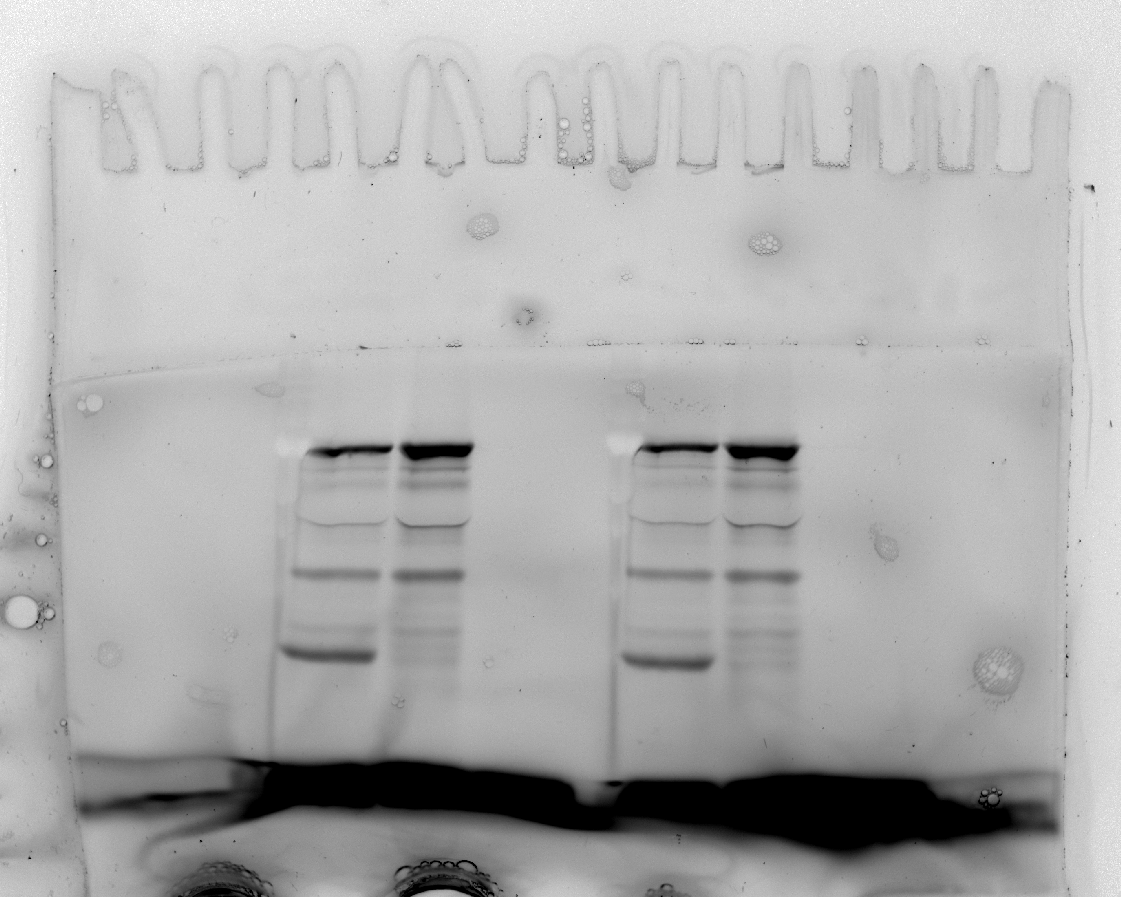

### Rep2 and Rep3 - ead11 - Laura Sierra 2024-12-06 14h56m31s(Composite).tif

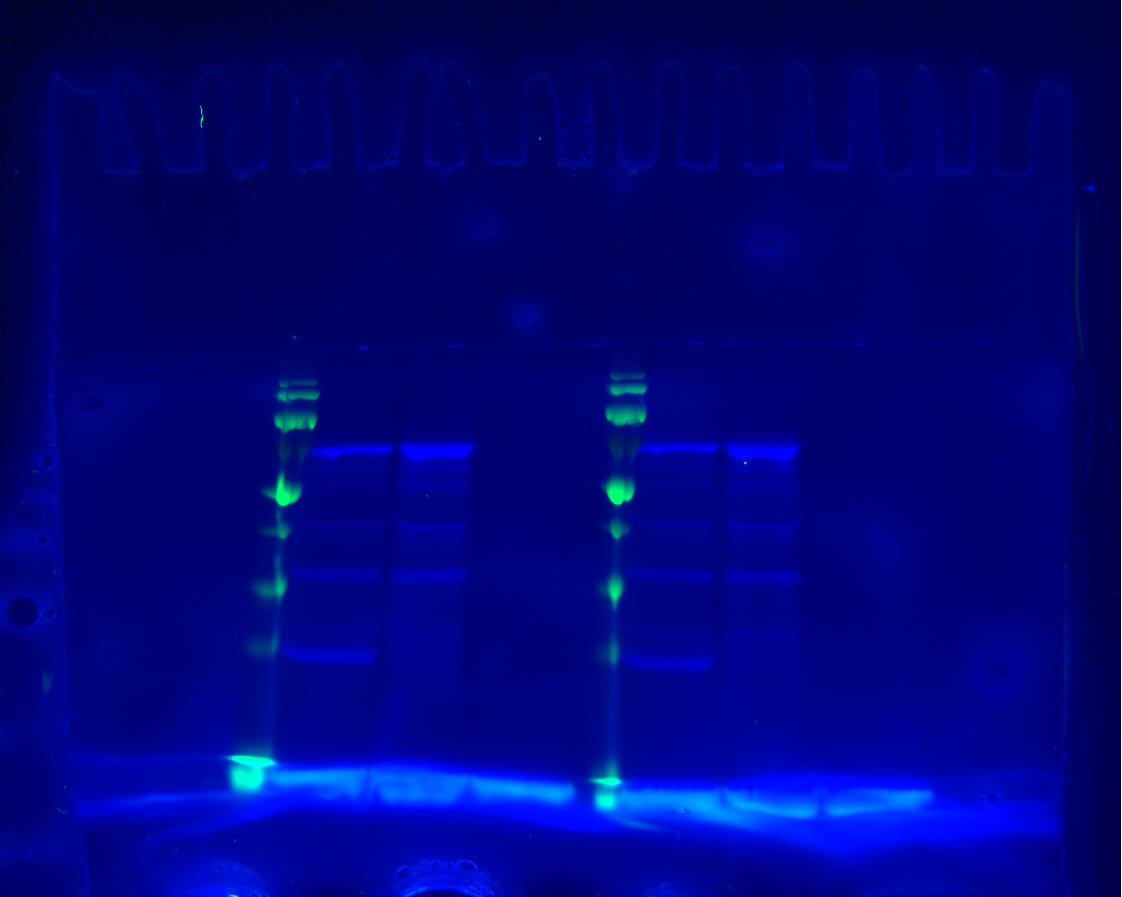

### Rep2_IMG20241015111310.jpg

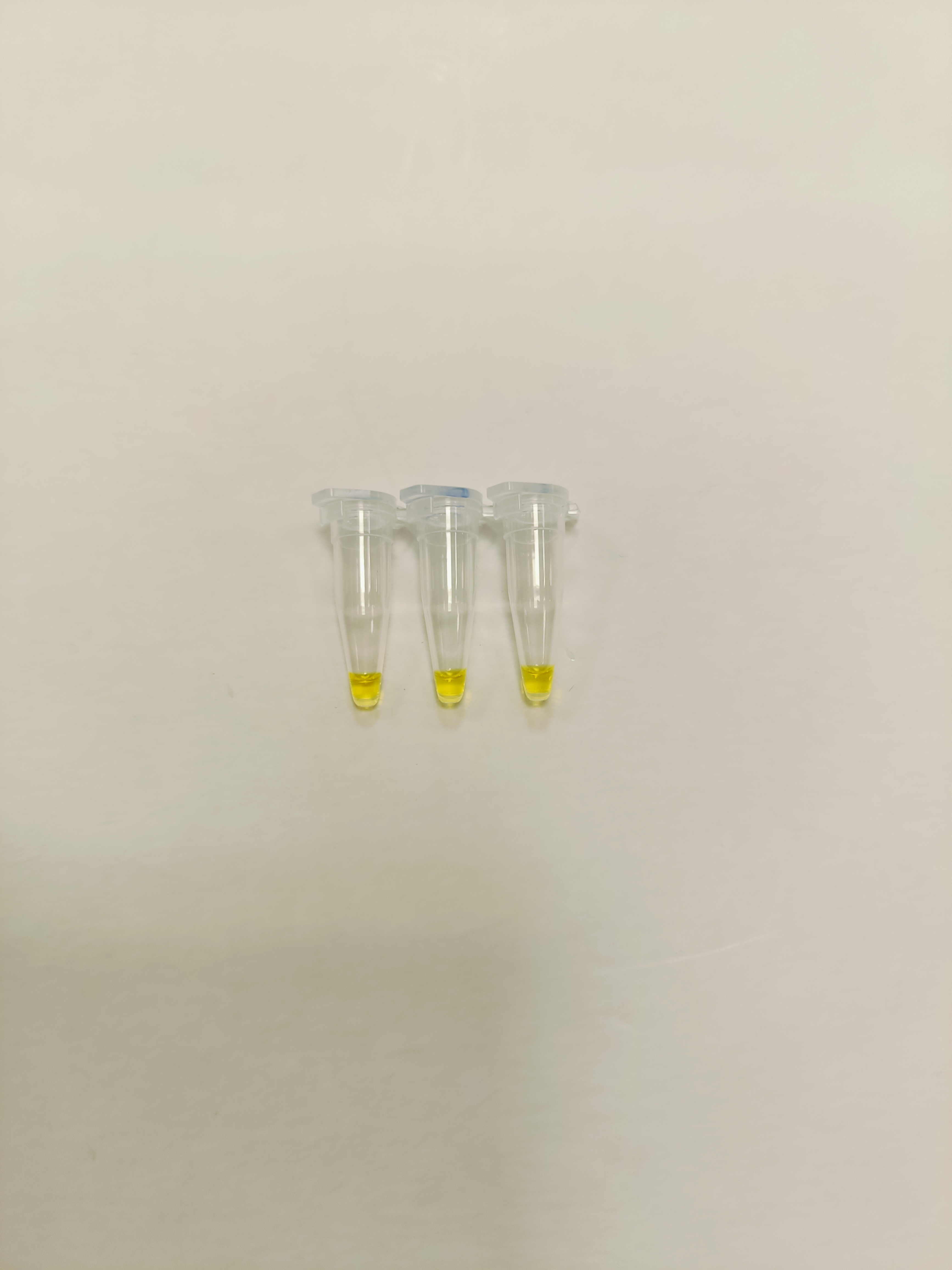

### Rep2_IMG20241015150924.jpg

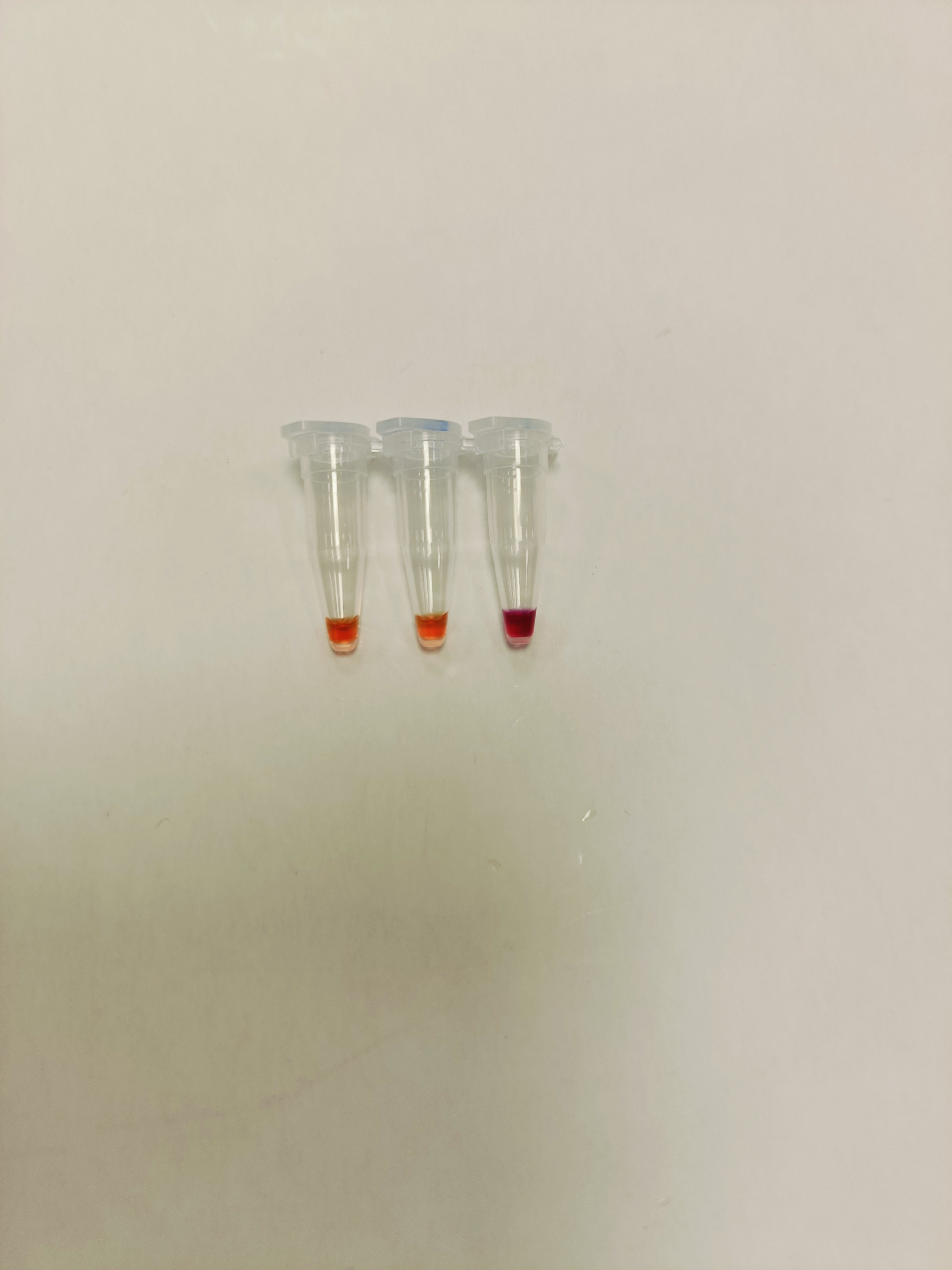

### Rep3 - ead11 - Laura Sierra 2024-10-04 09h19m26s(Alexa 488).tif

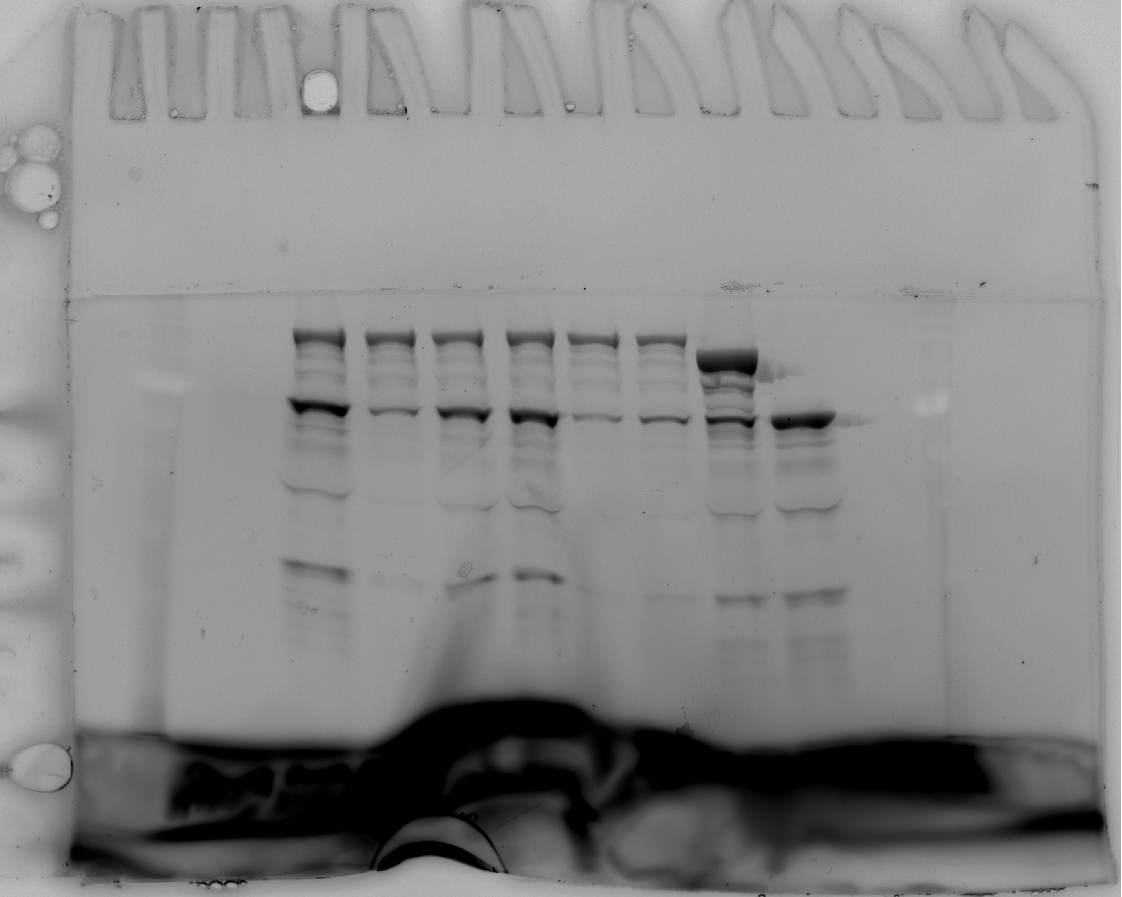

### Rep3 - ead11 - Laura Sierra 2024-10-04 09h19m26s(Composite).tif

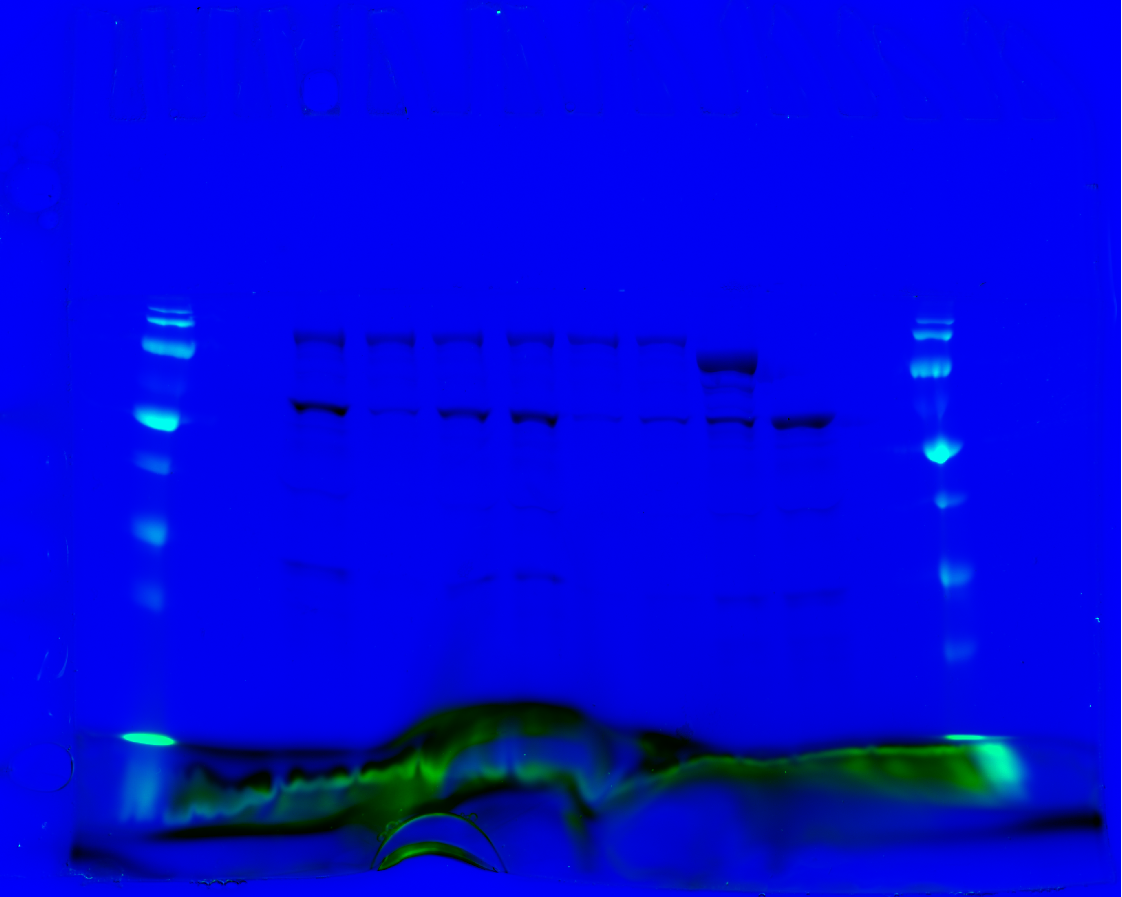

### Rep3_IMG20241015111322.jpg

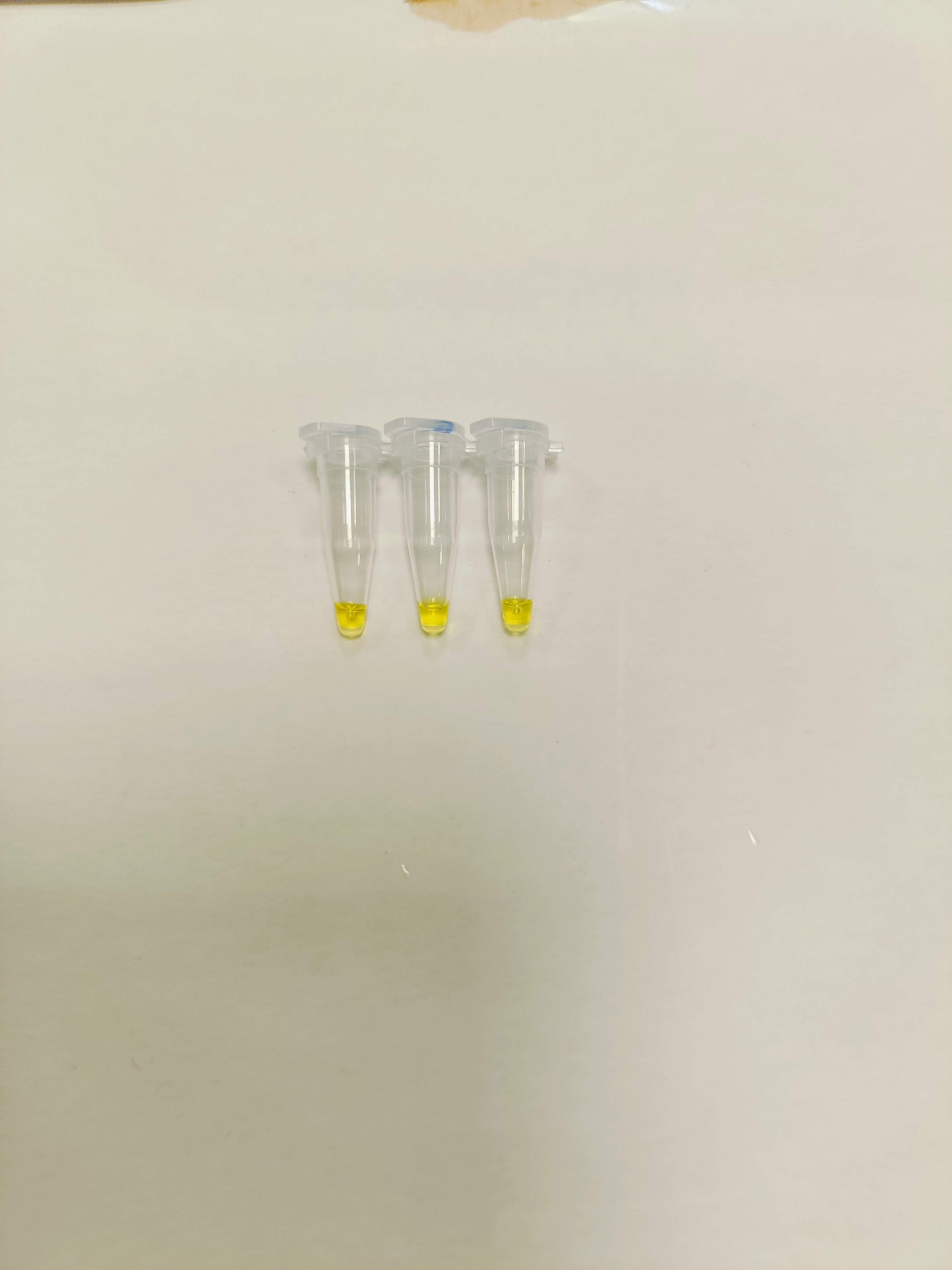

### Rep3_IMG20241015151137.jpg

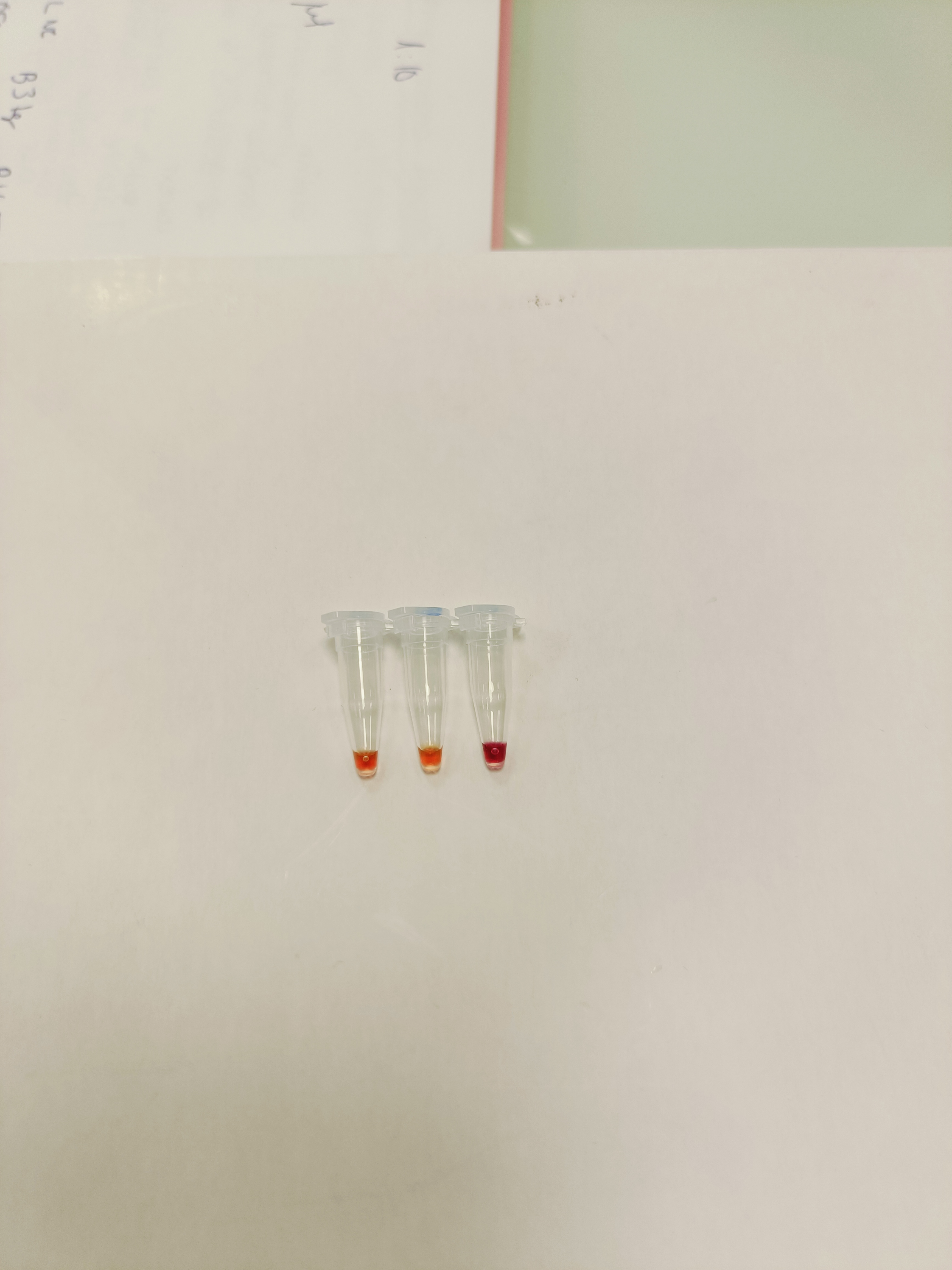

### Rep 1 - ead11 - Laura Sierra 2024-12-06 14h48m46s(Composite).tif

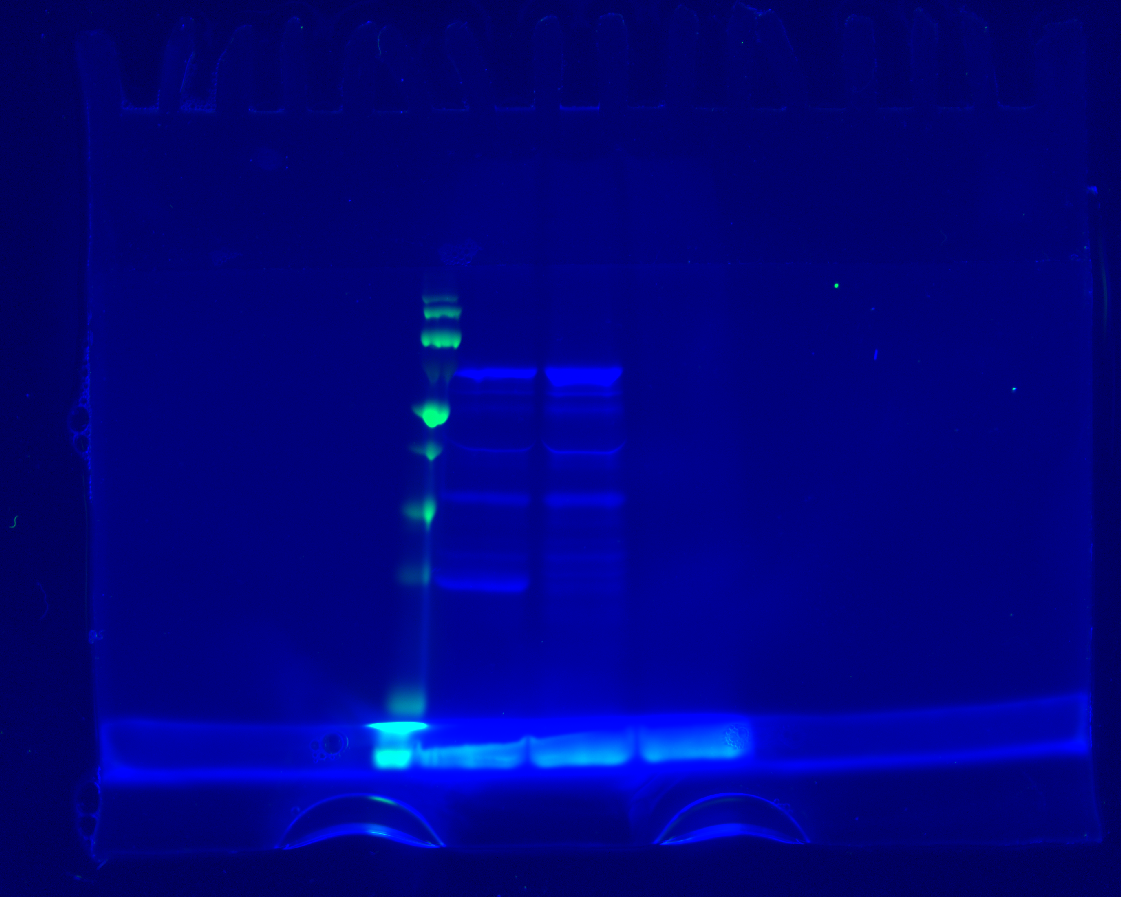
